# Unidirectional fork movement coupled with strand-specific histone incorporation ensures asymmetric histone inheritance

**DOI:** 10.1101/242768

**Authors:** Matthew Wooten, Jonathan Snedeker, Zehra Nizami, Xinxing Yang, Rajesh Ranjan, Elizabeth Urban, Jee Min Kim, Joseph Gall, Jie Xiao, Xin Chen

## Abstract

DNA replication establishes asymmetric epigenomes

**Summary:** One of the most fundamental questions in developmental biology concerns how cells with identical genomes differentiate into distinct cell types. One important context for understanding cell fate specification is asymmetric cell division, where the two daughter cells establish different cell fates following a single division. Many stem cells undergo asymmetric division to produce both a self-renewing stem cell and a differentiating daughter cell^1–5^. Here we show that histone H4 is inherited asymmetrically in asymmetrically dividing *Drosophila* male germline stem cells, similar to H3^6^. In contrast, both H2A and H2B are inherited symmetrically. By combining superresolution microscopy with the chromatin fiber method, we are able to study histone inheritance patterns on newly replicated chromatin fibers. Using this technique, we find asymmetric inheritance patterns for old and new H3, but symmetric inheritance patterns for old and new H2A on replicating sister chromatids. Furthermore, co-localization studies on isolated chromatin fibers and proximity ligation assays on intact nuclei reveal that old H3 are preferentially incorporated by the leading strand while newly synthesized H3 are enriched on the lagging strand. Finally, using a sequential nucleoside analog incorporation assay, we detect a high incidence of unidirectional DNA replication on germline-derived chromatin fibers and DNA fibers. The unidirectional fork movement coupled with the strand preference of histone incorporation could explain how old and new H3 are asymmetrically incorporated by replicating sister chromatids. In summary, our work demonstrates that the intrinsic asymmetries in DNA replication may help construct sister chromatids enriched with distinct populations of histones. Therefore, these results suggest unappreciated roles for DNA replication in asymmetrically dividing cells in multicellular organisms.

## Main Text

In the process of cell fate specification, epigenetic mechanisms play important roles by altering chromatin structure and gene expression patterns while preserving primary DNA sequences. Asymmetric cell division (ACD) has been characterized in multiple systems where it plays an essential role in generating cells with distinct fates in development, homeostasis, and tissue regeneration^3,4,7,8^. Stem cells, in particular, often use ACD to give rise to one daughter cell capable of self-renewal and another daughter cell in preparation for terminal differentiation^1–5^. In spite of the crucial role that epigenetic mechanisms play in regulating cell fate decisions during development^9–11^, it remains unclear how stem cells and differentiating daughter cells establish different epigenomes following ACD.

The *Drosophila* male germline stem cell (GSC) system provides a great model to investigate the fundamental molecular and cellular mechanisms underlying ACD^12,13^. Previous work has demonstrated that during the process of GSC ACD, old histone H3 are selectively segregated to the GSC, whereas new H3 are enriched in the gonialblast (GB) committed for differentiation^6,14^. These results indicate that stem cells may selectively retain preexisting histones that help define stem cell identity, whereas the differentiating daughter cell resets its epigenome as an initial step in the cellular differentiation program.

In eukaryotic cells, just as DNA must be duplicated *via* replication, chromatin must likewise be established on both strands during and after replication^15,16^. Accordingly, the bulk of canonical histones (i.e. H3, H4, H2A, and H2B) are synthesized and incorporated during DNA replication^17^. Old histones incorporated in nucleosomes on the parental DNA must be disassembled ahead of the replication fork and reassembled onto one of the two new double stranded DNA (dsDNA) templates behind the fork following DNA synthesis^18–20^. Although the process of new histone incorporation onto DNA has been well studied, how old histones are recycled during DNA replication is less clear^21–23^. Previous studies have shown that old histones can display a strand preference towards either the leading strand^24–29^, or the lagging strand^30,31^ during recycling events in different systems. Noticeably, the mode of histone incorporation has not been systematically studied in any multicellular organism in the context of cellular differentiation and asymmetric cell division. Furthermore, previous studies using biochemistry or high-throughput sequencing methods have not allowed for visualization of histone incorporation pattern at the single-molecule level. Characterizing patterns of histone incorporation during DNA replication in cells under their physiological condition is critical to our understanding of epigenetic regulation in animal development and various diseases, including cancer and tissue dystrophy [reviewed in^32^].

Using a heat shock-controlled switching system to label old histone with GFP (green fluorescent protein) and new histone with mKO (monomeric Kusabira Orange fluorescent protein) (Figures 1a and Supplementary figure 1a), we explored the inheritance pattern for all canonical histones following the ACD of *Drosophila* male GSCs. The distributions of old histone (GFP) and new histone (mKO) were measured following the second mitosis after heat shock-induced genetic switch^6^. Since mitotic GSCs account for less than 2% of the total population of GSCs^6,33–35^, post-mitotic GSC-GB pairs derived from the ACD of GSCs were used to visualize and quantify histone inheritance patterns in fixed cells (Supplementary Materials and Methods). For H4, we found that old H4-GFP was enriched in the GSCs on average 3.3-fold (Figures 1b and 1d), similar to what was previously reported for old H3^6,36^. By contrast, such an asymmetric old H4 inheritance pattern was not observed in S-phase spermatogonial (SG) pairs after symmetric cell division (Figures 1c and 1d). On the other hand, new H4-mKO displayed a more symmetric pattern between GSCs and GBs (Figures 1b and 1d). Presence of newly synthesized H4-mKO in both nuclei of the GSC-GB pair was consistent with the fact that both cells underwent S phase after the second mitosis following heat shock, which was further confirmed by the incorporation of the nucleoside analog EdU (5-ethynyl-2′-deoxyuridine) (Figures 1a and 1b). In mitotic GSCs at anaphase (Supplementary figure 1b) or telophase (Supplementary figure 1c), both old H4-GFP and new H4-mKO showed asymmetric segregation patterns. By contrast, using similar experimental strategies, old and new H2A (Figures 1e and 1g) as well as old and new H2B (Figures 1f and 1h) showed symmetric inheritance patterns in post-mitotic GSC-GB pairs. Further investigation of mitotic GSCs at anaphase confirmed this globally symmetric inheritance pattern for both H2A (Supplementary figure 1d) and H2B (Supplementary figure 1e). Additionally, both H2A (Figure 1g and Supplementary figure 1f) and H2B (Figure 1h and Supplementary figure 1g) displayed symmetric old and new histone inheritance patterns in post-mitotic SG pairs. Finally, the linker histone H1 also showed globally symmetric inheritance pattern in post-mitotic GSC-GB pairs (Supplementary figure 1h).

**Figure 1:**
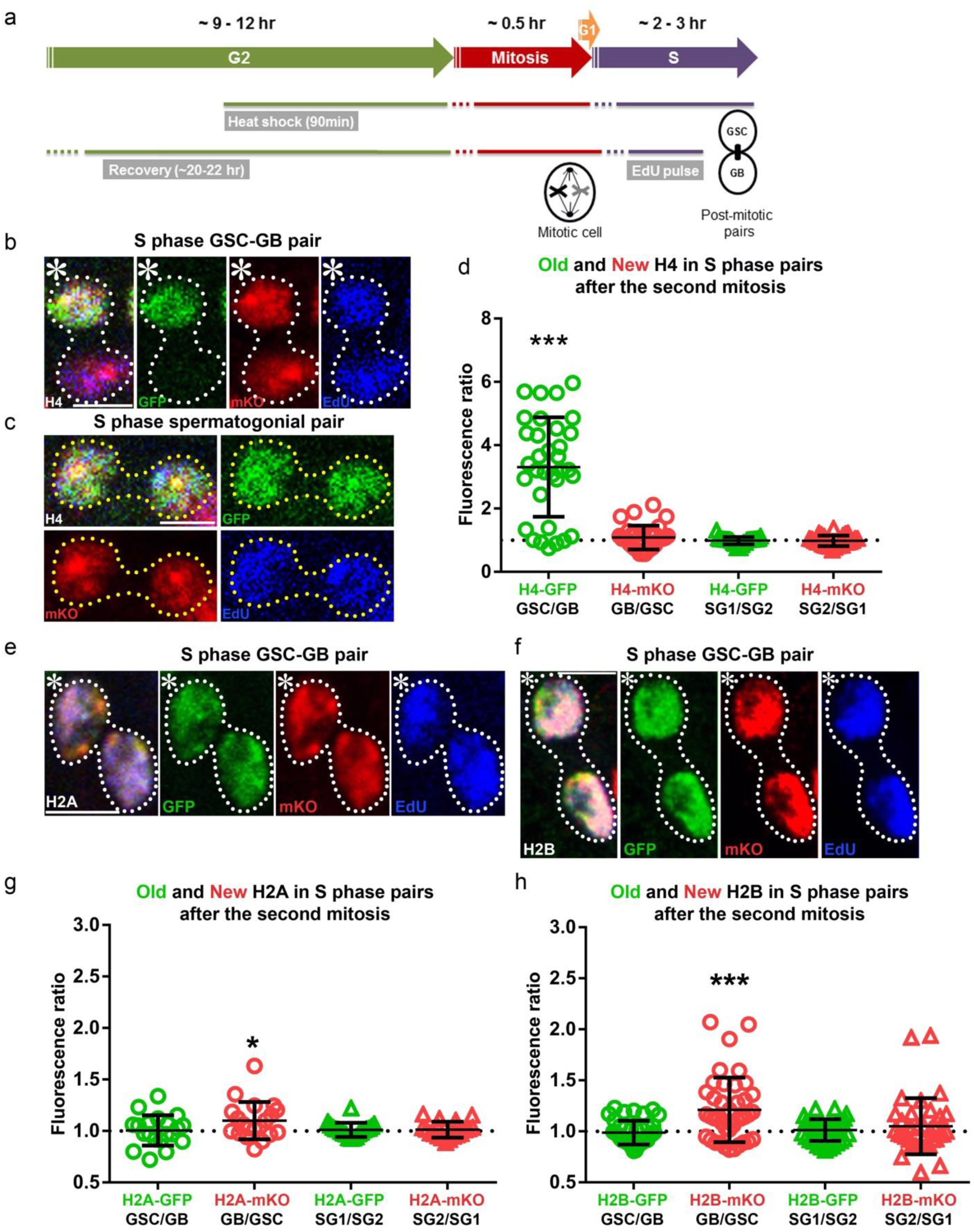
Histone H4 shows asymmetric while histones H2A and H2B show symmetric inheritance patterns during *Drosophila* GSC asymmetric divisions. **(a)** A cartoon depicting the experimental design. (**b**) H4 distribution patterns in a post-mitotic GSC-GB pair labeled with EdU (blue): H4-GFP (green) is distributed asymmetrically towards the GSC whereas H4-mKO (red) distributed more evenly between the GSC and the GB. (**c**) H4 distribution patterns in a post-mitotic SG pairs. Both H4-GFP and H4-mKO are symmetrically distributed between the two SG nuclei. (**d**) Quantification of both H4-GFP and H4-mKO distributions in GSC-GB pairs (*n*=33) and SG1-SG2 pairs (*n*=27). See Supplementary table 1 for details. (**e**) Symmetric H2A inheritance pattern in a post-mitotic GSC-GB pair. **(f)** Symmetric H2B inheritance pattern in a post-mitotic GSC-GB pair. (**g**) Quantification of H2A-GFP and H2A-mKO distribution in GSC-GB pairs (*n*=20) and SG1-SG2 pairs (*n*=20). See Supplementary table 2 for details. (**h**) Quantification of H2B-GFP and H2B-mKO distribution in GSC-GB pairs (*n*=40) and SG1-SG2 pairs (*n*=36). See Supplementary table 3 for details. *** *P* < 0.0001, * *P* < 0.05, two-tailed student’s *t*-test if average significantly different than 1. Error bars represent 95% confidence interval. Both new H2A and new H2B show a subtle, but statistically significant enrichment in GB compared to GSC in post-mitotic pairs, likely due to asynchronous ongoing S phase in both GB and GSC nuclei. Scale bar: 5μm. Asterisk: hub.

These findings indicate that even though H3, H4, H2A, and H2B are all initially incorporated in a replication-dependent manner, different histones display distinct inheritance patterns in *Drosophila* male GSCs. In order to directly examine histone incorporation patterns on newly replicated DNA, we adapted the chromatin fiber technique^37–39^ to visualize EdU pulse-labeled DNA with associated proteins outside the confines of the nucleus (Materials and Methods). To validate this technology, chromatin fibers were isolated from *Drosophila* embryos at the syncytial blastoderm stage and compared with previous electron microscopy images^18^. Replicating regions (EdU-positive) of chromatin fibers ranged from 250 nm to 8μm long, similar in size to replicating regions identified with electron microscopy ^18,40,41^. EdU incorporation clearly distinguished unreplicated (Supplementary figures 2a and 2b, 2j and 2k) and newly replicated (Supplementary figures 2a and 2c, 2j and 2l) regions of chromatin fibers. Consistent with prior findings^37^, EdU-positive regions showed wider fiber structure and brighter DNA staining with DAPI (Supplemental figures 2a and 2d) or YOYO-3 (Supplemental figure 2m) DNA dye.

To confirm that DAPI-bright, EdU-positive fiber structures represent replicating regions, fibers were isolated from non-replicating cells (i.e. *Drosophila* adult eye) treated with EdU. Fibers isolated from non-replicating cells showed uniform DAPI staining with no identifiable regions of EdU incorporation (Supplemental figures 2f-i) compared to those derived from replicating cells (Supplemental figures 2a-e). These data demonstrate that DAPI-bright, EdU-positive chromatin fibers represent regions of DNA synthesis.

Using confocal microscopy, only 3.2% (*n*=250) of DAPI-bright, EdU-positive regions on embryo-derived chromatin fibers could be clearly resolved into two sister chromatids (Supplemental figures 2j and 2l). To overcome resolution limits, we used two high resolution microscopy methods: Stimulated Emission-Depletion (STED) microscopy and Zeiss Airyscan imaging. Both STED (Supplementary figures 2m and 2o; Supplementary figure 3a and 3b) and Airyscan (Supplementary figure 3c and 3d) greatly improved the frequency of resolving sister chromatids at actively replicating regions of chromatin fibers. Overall, the percentage of spatially resolvable sister chromatids from EdU-positive chromatin fibers ranged from 8.6% using Airyscan to 25.0% using STED (Supplementary figure 2p). Differences in the relative frequency of resolvable sisters between these two methods likely reflects the lower resolution of Airyscan (∼150 nm)^42^ compared to STED (∼35 nm)^43^. The application of superresolution microscopy to imaging replicating chromatin fibers represents a new methodology to study DNA replication and nucleosome assembly.

Using this method, we next explored old and new histone distribution on chromatin fibers isolated from the early-stage *Drosophila* male germ cells (Materials and Methods). Using Airyscan imaging, unreplicated EdU-negative regions were detected as a single fiber structure enriched with predominantly old histones (Figures 2a, 2b, 2h, and 2i). To explore histone incorporation patterns on newly replicated chromatin fibers, we compared the distribution of old and new H2A and H3 on sister chromatids. Old and new H2A showed a largely symmetric distribution on chromatin fibers (Figures 2a and 2c). By contrast, old and new H3 showed a more asymmetric distribution pattern on newly replicated sister chromatids (Figures 2h and 2j). These results with H3 were confirmed using two-color STED imaging (Figures 2k and 2m).

**Figure 2:**
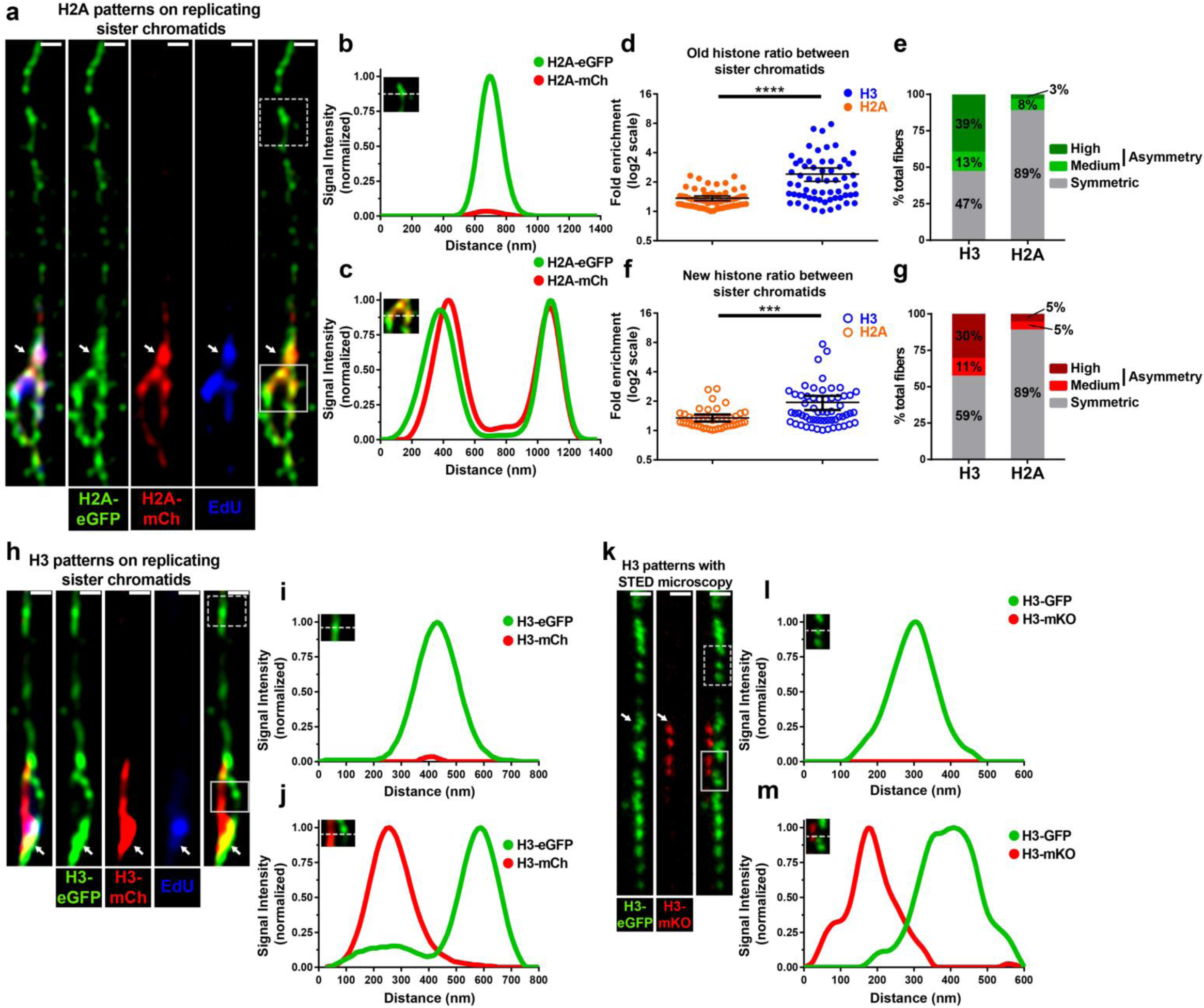
Asymmetric H3 and symmetric H2A distribution on replicating sister chromatids. **(a)** Airyscan image of chromatin fiber labelled with EdU showing old H2A-eGFP (green) and new H2A-mCherry (red) distribution on unreplicated and replicating regions. At the replicating region with EdU and new H2A signals, sister chromatids show symmetric old and new H2A distribution. Scale bar: 500 nm. (**b**) Line-plot shows old H2A-eGFP and new H2A- mCherry distribution on unreplicated region without EdU (box with dotted white lines in **a**, inset in **b**). (**c**) Line-plot shows old H2A-eGFP and new H2A-mCherry distribution on replicated region with both EdU and new H2A (box with solid white lines in **a**, inset in **c**). (**d**) Quantification of old H2A (avg. 1.36; *n*=65 replicating regions; *n =* 33 chromatin fibers) and old H3 (avg. 2.41; *n*=61 replicating regions; *n =* 35 chromatin fibers) distribution between sister chromatids at replication regions on chromatin fibers. (**e**) Classification of old histone patterns H3 *versus* H2A. We classified sister chromatids as symmetric (ratio <1.8-fold difference), moderately asymmetric (1.8 < ratio < 2.44) or highly asymmetric (ratio > 2.44-fold difference). We used 2.44 as a standard for calling fibers highly asymmetric, as it is two standard deviations above the average ratio observed between old H2A fibers. (**f**) Quantification of new H2A (avg. 1.24; *n*=45 replicating regions; 30 chromatin fibers) and new H3 (avg. 1.94; *n*=59 replicating regions; 32 chromatin fibers) distribution between sister chromatids at replication regions on chromatin fibers. **** *P* < 0.0001, *** *P* < 0.001, Mann-Whitney U test. Error bars represent 95% confidence interval. (**g**) Classification of new histone patterns H3 *versus* H2A. We classified sister chromatids as symmetric (ratio <1.70-fold difference), moderately asymmetric (1.8 < ratio < 2.16) or highly asymmetric (ratio > 2.16-fold difference). We used 2.16 as a standard for calling fibers highly asymmetric, as it is two standard deviations above the average ratio observed between old H2A fibers. (**h**) Airyscan image of chromatin fiber labelled with EdU showing old H3-eGFP (green) and new H3-mCherry (red) distribution on unreplicated and replicating regions. At the replicating region with EdU and new H3 signals, sister chromatids show asymmetric old and new H3 distribution. Scale bar: 500 nm. (**i**) Line-plot shows old H3- eGFP and new H3-mCherry distribution on unreplicated region without either EdU or new H3 (box with dotted white lines in **h**, inset in **i**). (**j**) Line-plot shows old H3-eGFP and new H3- mCherry distribution on replicated region with both EdU and new H3 (box with solid white lines in **h**, inset in **j**). (**k**) Two-color STED image of chromatin fiber showing old H3-GFP and new H3-mKO distribution on unreplicated and replicating chromatin region. New H3 incorporation is confined to regions that show double fiber structure associated with active DNA replication. The transition from single fiber to double fiber occurs at the point where new histone incorporation begins (white arrow). (**l**) Line-plot shows old H3-GFP and new H3-mKO distribution on unreplicated region without new H3 (box with dotted white lines in **k**, inset in **l**). (**m**) Line-plot shows old H3-GFP and new H3-mKO distribution on replicated region with new H3 (box with solid white lines in **k**, inset in **m**).

To systematically compare histone distribution patterns of H3 and H2A along sister chromatids, we divided sister chromatid fibers into 2μm units and measured the fluorescence levels for both old and new histones on each unit (Supplementary figure 3e, Materials and Methods). Overall, old H3 displayed a significantly higher incidence of asymmetry than did H2A fibers (Figure 2d, *P* < 10^−4^). Old H3 showed on average a 2.41-fold ratio while old H2A showed a 1.36-fold ratio between sister chromatids. Additionally, new H3 also displayed a significantly higher incidence of asymmetry when compared to new H2A (Figure 2f, *P* < 10^−3^). New H3 showed on average a 1.94-fold difference while new H2A showed a 1.24-fold difference between sister chromatids.

To further understand the differences in old and new histone incorporation patterns between H3 and H2A, we classified fibers as symmetric, moderately asymmetric or highly asymmetric (Materials and Methods). Using these criteria, 39% of H3 fibers were found to be highly asymmetric compared to just 3% of H2A fibers for old histones (Figure 2e, *P* < 10^−4^). Similarly, 30% of H3 fibers were found to be highly asymmetric compared to just 5% of H2A fibers for new histones. For the moderately asymmetric fibers, H3 and H2A fibers showed comparable frequencies: 13% of H3 fibers were moderately asymmetric compared to 8% of H2A fibers for old histones (Figure 2e, *P*=0.32); and 11% of H3 fibers showed moderate asymmetry compared to 5% for H2A for new histones (Figure 2g, *P*=0.33). In summary, these results demonstrate that both old and new H3 are more asymmetrically incorporated during DNA replication compared to old and new H2A, consistent with their distinct segregation patterns during ACD of GSCs (Figure 1 and Supplementary fig. 1).

As old and new H3 show significant asymmetries during the process of replication-coupled nucleosome assembly, we next tested whether old *versus* new H3 asymmetry correlates in any way with strand-enriched DNA replication machinery components. To determine strand specificity, chromatin fibers were isolated from flies expressing eGFP-RPA (replication protein-A) fusion protein under the control of the endogenous regulatory elements of the *rpa* gene (*rpa>RPA-eGFP)*^44^. RPA represents a highly conserved single-stranded DNA-binding protein significantly enriched at the lagging strand^45^. To visualize old histones, we utilized an antibody against the H3K27me3 histone modification, which has been shown to be enriched on old H3^46^. At EdU-positive regions where the sister chromatids could be resolved, RPA and H3K27me3 occupied opposite strands of the bubble structure (Figures 3a and 3b), suggesting that old H3 is recycled to the leading strand. Quantification showed an average of 3.2-fold more H3K27me3 at the RPA-depleted leading strand compared to the RPA-enriched lagging strand (Figure 3c, *P* < 10^−4^.). Furthermore, when all fibers were classified into three categories: leading strand enriched, symmetric, or lagging-strand enriched, 64% of fibers showed leading-strand enrichment, 30% of fibers were symmetric and only 6% of fibers showed lagging strand enrichment (Figure 3d). We also investigated H4 incorporation patterns at replicating regions using an old H4-enriched H4K20me2/3 modification^46,47^. At EdU-positive regions of germline-derived chromatin fibers, H4K20me2/3 levels were more abundant on the RPA-negative leading strand when compared to the RPA-positive lagging strand (Supplementary figures 4a and 4b). Quantification showed a 2.1-fold difference on average in H4K20me2/3 levels of leading strand compared to the lagging strand (Supplementary figures 4c, *P* < 10^−4^). Further analysis demonstrated that 54% of fibers showed old histone enrichment towards the leading strand, 31% showed symmetry, while only 15% of fibers showed enrichment towards the lagging-strand (Supplementary figure 4d). Taken together, these results suggest that old H4, similar to old H3, is preferentially recycled to the leading strand.

**Figure 3:**
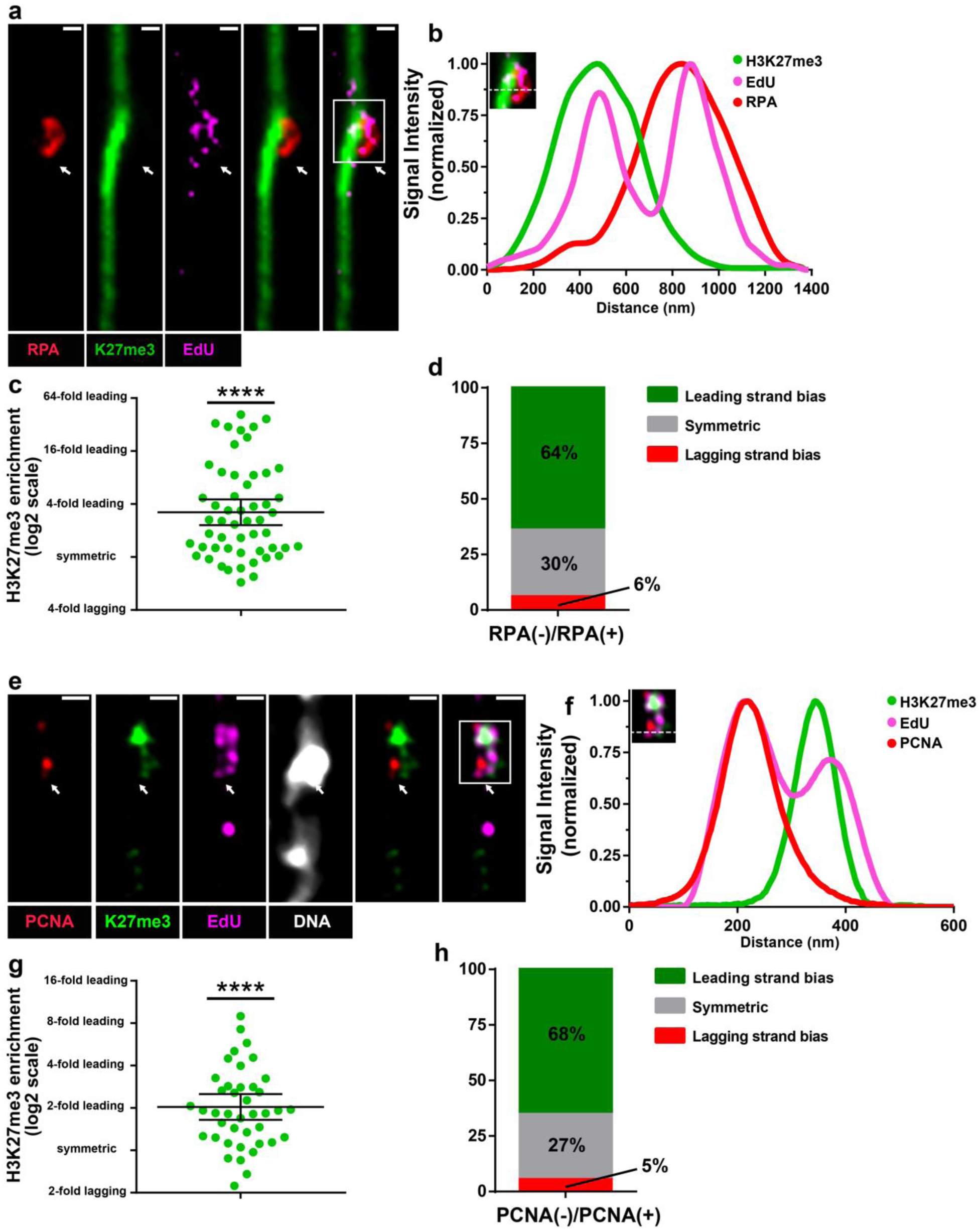
Old H3 preferentially associate with the leading strand on chromatin fibers. **(a)** Confocal image of chromatin fiber labelled with EdU showing anti-correlated H3K27me3 and RPA distribution. The transition from single fiber to double fibers is correlated with the EdU incorporation site (white arrow). (**b**) Line-plot shows EdU, H3K27me3 and RPA distribution across the replicating region (box with solid white lines in **a**, inset in **b**). (**c**) Quantification of the log2 (average H3K27me3 fluorescence intensity on RPA-depleted sister chromatid/ average H3K27me3 fluorescence intensity on RPA-enriched sister chromatid) (Average fold enrichment=3.20-fold, *n*=53 replicating regions from 35 chromatin fibers). Y-axis is with log2 scale. Data is significantly different from 0. **** *P* < 0.0001, Mann-Whitney U test. (**d**) Classification of RPA-labeled sister chromatids into leading-strand enriched (ratio >1.4), lagging-strand enriched (ratio <1.4) and symmetric (−1.4 < ratio < 1.4). (**e**) Airyscan image of chromatin fiber labelled with EdU showing anti-correlated H3K27me3 and PCNA distribution. The white arrow indicates the replication bubble. (**f**) Line-plot shows EdU, H3K27me3 and PCNA distribution across replicating region (box with solid white lines in **e**, inset in **f**). (**g**) Quantification of the log_2_ (average H3K27me3 fluorescence intensity on PCNA-depleted sister chromatid/ average H3K27me3 fluorescence intensity on PCNA-enriched sister chromatid) (Average fold enrichment 2.04; *n*=41 replicating regions from 34 chromatin fibers). Y-axis is with log2 scale. Data is significantly different from 0. **** *P* < 0.0001, Mann-Whitney U test (*n*=52). (**h**) Classification of PCNA-labelled fibers into leading-strand enriched (inter-sister ratio >1.4), lagging-strand enriched (inter-sister ratio <1.4) and symmetric (−1.4 < inter-sister ratio < 1.4). Y-axis is with log2 scale. **** *P* < 0.0001, Mann-Whitney U test. Error bars represent 95% confidence interval in (**c**) and (**g**).

To further validate histone inheritance patterns at the replication fork, similar experiments were performed using another lagging-strand-enriched component, PCNA (Proliferating Cell Nuclear Antigen)^45^, which was expressed in its endogenous genomic context (*pcna>PCNA-eGFP)*^44^. At EdU-positive sister chromatid regions, PCNA and H3K27me3 occupied opposite sister chromatids (Figures 3e and 3f), further demonstrating that old H3 is preferentially recycled to the leading strand. Quantification showed an average of 2.0-fold more H3K27me3 at the PCNA-depleted leading strand compared to the PCNA-enriched lagging strand (Figure 3g, *P* < 10^−4^). Further analysis showed 68% of fibers showed old histone enrichment towards the leading strand, 27% showed symmetry, while only 5% of fibers showed enrichment towards the lagging-strand (Figure 3h). Taken together, these results demonstrate that during DNA replication, old (H3-H4)_2_ tetramers are preferentially recycled by the leading strand.

Previous studies have shown that asymmetric histone inheritance is specific to asymmetrically dividing GSCs but not to late-stage SGs^6^. To investigate whether this cellular specificity originates from histone incorporation differences at the replication fork, we utilized a late-stage germ cell driver *bam-Gal4* to exclusively express the H2A-GFP in SGs to label late-stage germ cell-derived chromatin fibers. The EdU-positive and H2A-GFP-labeled fibers were derived from SGs whereas EdU-positive but H2A-GFP-negative fibers likely came from early-stage germ cells including GSCs. Using H3K27me3 as a proxy for old H3, chromatin fibers from early-stage germ cells (Supplementary figures 5a-b) showed higher frequency and more substantial asymmetry between sister chromatids compared to fibers derived from late-stage SGs (Supplementary figure 5d-f). Early-stage germ cells showed a 2.5-fold difference between sister chromatids whereas late-stage germ cells showed only a 1.6-fold difference (Supplementary figures 5g and 5h, *P* < 10^−3^). These findings suggest that asymmetries in histone inheritance decrease as germ cells begin to differentiate.

As a complementary method to explore histone inheritance patterns at the replication fork in intact nuclei, we used an imaging based proximity ligation assay (PLA) to probe the spatial proximity between histones (old *versus* new) and different strand-enriched DNA replication components. We used CRISPR/Cas9-mediated genome editing^48,49^ to tag the lagging strand-enriched DNA ligase at its endogenous genomic locus using a 3xHA epitope. We then applied anti-HA for the PLA assay to probe the spatial proximity between DNA ligase and old *versus* new histones. We observed a higher number of PLA fluorescent puncta between ligase and new H3-mKO (Supplementary figure 6a) than those between ligase and old H3-GFP (Supplementary figure 6b). Quantification of the overall PLA signals in GSCs showed significantly more PLA fluorescent puncta between ligase and new H3 than those between ligase and old H3 (Supplementary figure 6c, *P*<0.01). Using another lagging strand-enriched component, PCNA, as a marker for the PLA experiments, we also observed a higher number of PLA fluorescent puncta between PCNA and new H3-mKO (Supplementary figure 6d) than those between PCNA and old H3-GFP (Supplementary figure 6e). Again, quantification of the overall PLA signals showed significantly more PLA fluorescent puncta between PCNA and new H3-mKO than those between PCNA and old H3-GFP in GSCs (Supplementary figure 6f, *P*<10^−3^). As a control, we also performed PLA experiments using a strain where the tags for old H3 and new H3 were swapped, resulting in old H3-mKO and new H3-GFP. Consistent with the previous results, more PLA fluorescent puncta were obtained between PCNA and new H3-GFP (Supplementary figure 6g) than the signals between PCNA and old H3-mKO (Supplementary figure 6h; quantified in supplementary figure 6i, *P*<0.05). To confirm the specificity of our PLA signal, we performed PLA in non-replicating somatic hub cells as well as between histones and a cytoplasmic protein Vasa^50^ (Supplementary figure 6j). In these experiments, we observed negligible PLA signal, confirming that PLA signals were specific to replicating nuclei and false positive signals were minimal in our experimental conditions. Together, these results are consistent with the chromatin fiber results shown above (Figure 3), suggesting that new H3 preferentially associates with the lagging strand.

Unlike GSCs, SGs showed no significant strand preference between old and new histones for either ligase (Supplementary figure 6c) or PCNA (Supplementary figure 6f). Together, results from both chromatin fiber technique (Supplementary figure 5) and PLA method (Supplementary figure 6) demonstrate that histone distribution patterns show a cellular specificity not only during mitosis^6^ (Figures 1 and Supplementary figure 1), but also during DNA replication. We therefore conclude that differences in epigenetic inheritance at the replication fork likely underlie differences in global epigenetic inheritance patterns observed between GSCs and SGs.

Up to now, we have demonstrated that old H3 are incorporated on the leading strand whereas new H3 are preferentially incorporated on the lagging strand during the process of replication-coupled nucleosome assembly. However, if replication forks are proceeding outward from replication origins in a bidirectional manner, asymmetries in histone inheritance at the replication fork alone would lead to alternating stretches of leading-strand-incorporated old histones and lagging-strand-incorporated new histones on each of the two duplicating sister chromatids (Figure 4a), which would not be sufficient to explain the global asymmetry of histone inheritance we have observed. Therefore, we hypothesize that replication forks are coordinated to achieve long-range asymmetric histone patterns.

**Figure 4:**
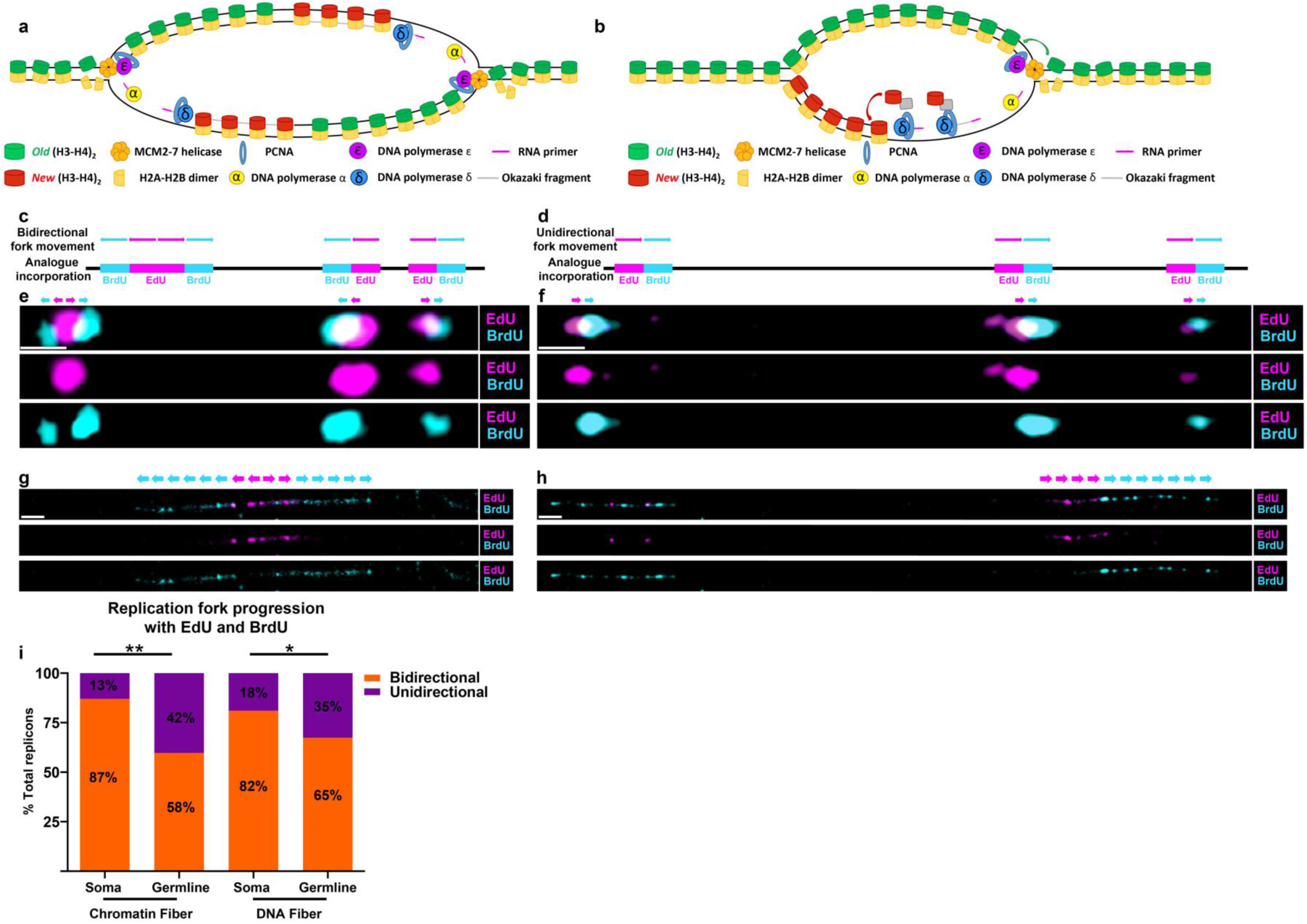
Germline-derived chromatin and DNA fibers show more unidirectional fork progression compared to soma-derived chromatin and DNA fibers. **(a)** A cartoon showing strand biased histone incorporation at a bidirectional replication fork. (**b**) A cartoon showing strand biased histone incorporation at a unidirectional replication fork. (**c**) Predicted bidirectional fork progression result. (**d**) Predicted unidirectional fork progression result. (**e**) Bidirectional fork progression pattern from somatic cell derived chromatin fiber. Replicons show early label (EdU in magenta) flanked by late label (BrdU in cyan) on both sides. (**f**) Unidirectional fork progression pattern from germline-derived chromatin fiber. Multiple replicons show alternation between early label (EdU in magenta) and late label (BrdU in cyan) along the chromatin fiber toward the same direction. (**g**) Bidirectional fork progression pattern from somatic cell derived DNA fiber. (**h**) Unidirectional fork progression pattern from germline-derived DNA fiber. (**i**) Quantification of fork progression patterns in somatic cell-derived *versus* germline-derived chromatin and DNA fibers. Germline-derived fibers show a significantly higher incidence of unidirectional fork progression: 42% in germline chromatin fiber (*n*=54) *versus* 13% in soma chromatin fiber (*n*=31), 35% in germline DNA fiber (*n*=109), 18% soma DNA fiber (*n*=45). **: *P* < 0.01,*: *P* < 0.05, Chi-squared test. Scale bar: 1μm for (**e**) and (**f**), 2μm for (**g**) and (**h**).

To explore the fork movement in the *Drosophila* germline, we applied a sequential nucleoside analog incorporation to the chromatin fiber method: active DNA replication regions were first labeled with EdU and subsequently by BrdU. A DNA dye was added to verify fiber continuity (Materials and Methods). Only continuous fibers containing multiple replicons in tandem were included for fork movement analysis (Supplemental figures 7a-7d). The progression of replication forks in a bidirectional (Figure 4c) or a unidirectional manner (Figure 4d) produces distinct patterns. Chromatin fibers derived from somatic cells, such as larval imaginal disk cells, displayed largely bidirectional fork movement (Figure 4e), as 87% of replicons on chromatin fibers showed typical bidirectional fork movement while only 13% of replicons showed unidirectional fork movement (*n*=31, Figures 4e). In contrast, a substantial fraction (42%) of germline-derived chromatin fibers contained replicons with unidirectional replication progression (*n*=53, Figures 4f). Furthermore, fork movement in unidirectional replicons appeared to be coordinated, as multiple unidirectional forks appeared to move in the same direction (Figure 4f). To further explore replication patterns in the *Drosophila* male germline, we utilized a similar sequential nucleoside analog incorporation with DNA fibers. Consistent with the chromatin fiber data, DNA fibers derived from adult testis showed higher incidence of unidirectional fork movement (35%, *n*=109, Figures 4h-i) compared to those derived from somatic tissues (18%, *n*=45, Figures 4g and 4i).

In summary, results using both chromatin fiber and DNA fiber methods reveal that replication could be coordinated in the *Drosophila* germline to allow for unidirectional fork movement (Figure 4b). Together with the detected strand bias found between old and new histones, these mechanisms could expand asymmetric histone incorporation at individual forks to global asymmetries between sister chromatids. While asymmetries in the deposition of histone proteins have been observed experimentally^19,24–28^, a majority of studies have demonstrated that on a global scale, old and new histones are equally associated with the leading and lagging strands following replication^15,51–53^. However, this question had not been addressed in a multicellular organism in a developmental context. Interestingly, recent studies on histone segregation in mouse embryonic stem cells^47^ and yeast^30^ reveal molecular mechanisms that act at the replication fork to counteract asymmetric histone incorporation, in order to achieve a more symmetric outcome. By contrast, studies in our system suggest that in a developmental context, asymmetries at the fork can be utilized as a tool to generate inheritable epigenetically distinct sister chromatids with crucial roles in regulating cell fate decisions^36^.

Unidirectional replication and coordinated fork movement are by no means unprecedented observations^54^. Fork block systems in the replicating *S. pombe* are utilized during mating-type switching to help coordinate fork movement across the mating type locus to create a DNA lesion necessary for initiating the DNA repair mechanisms involved in the process of mating-type switching^55^. In *Drosophila*, it has been shown that fork movement at the rDNA region is unidirectional^56^. Fork block systems have also been found in metazoan systems to ensure that replication/transcription collisions do not occur in the context of the heavily transcribed loci^57^. A majority of studies on mammalian replicons have identified that approximately 5-14% of origins are replicated in a unidirectional manner whereas 86-95% are bidirectional^58,59^. Some studies have observed that higher incidences of unidirectional fork movement can be detected in late-replicating regions of the genome when compared to early replicating regions^60^. However, fork coordination across broad stretches of the genome as a means to regulate epigenetic inheritance represents a previously uncharacterized aspect of cell-type specific regulation of DNA replication.

In this work, we have also optimized the chromatin fiber technique and combined it with different high spatial resolution microscopy methods to visualize sister chromatids as they are undergoing the processes of DNA replication and replication-coupled nucleosome assembly. Combining this technique with the dual-color histone labeling system and/or immunostaining for histone modifications, we have established a novel method to study replication-coupled nucleosome assembly at sister chromatids. Noticeably, this information would be very difficult to attain by other means. For example, any sequencing-based method would not be able to distinguish sister chromatids where origins of replications are not well characterized, which is the case for most multicellular organisms.

Together our findings suggest that DNA replication may play a novel, unappreciated role in directing histone incorporation to differentially establish epigenetic information on two genetically identical sister chromatids. Furthermore, these results identify that DNA replication can be exploited in a cell-type specific manner. While the molecular players responsible for this cell-type-specificity remain unclear, this demonstration of a potential regulatory role for DNA replication represents an important step forward in understanding how DNA replication and replication-coupled nucleosome assembly regulate asymmetric cell division and cell fate specification.

## Author contributions

Conceptualization, M.W., Z.N., X.Y., J.S., J.G., J.X. and X.C.; Methodology, M.W., Z.N., X.Y., J.S., J.G., J.X., and X.C.; Investigation, M.W., Z.N., R.R., J.S., J-M.K., E.U.; Writing – Original Draft, M.W., Z.N., X.Y., J.S., J.G., J.X. and X.C.; Funding Acquisition, J.X., J.G. and X.C.; Supervision, J.X., J.G. and X.C.

## Acknowledgements

We thank B. Shelby and E. Wieschaus for the RPA-GFP fly line. We thank E. Moudrianakis, A. Spradling, J. Berger, M. Van Doren, R. Johnston and X.C. lab members for suggestions. We thank Johns Hopkins Integrated Imaging Center for confocal imaging and Carnegie Institute Imaging Center for STED microscopy work. Supported by NIH 5T32GM007231 and F31GM115149-01A1 (M.W.), NIH R01GM112008 (J.X.), NIH R01GM33397 (J.G.), NIH R01GM112008, R35GM127075, the Howard Hughes Medical Institute, the David and Lucile Packard Foundation, and Johns Hopkins University startup funds (X.C.)

## Materials & Correspondence

Correspondence and material requests should be addressed to X.C.

## REFERENCES

1. Knoblich, J.A. Asymmetric cell division: recent developments and their implications for tumour biology. Nat Rev Mol Cell Biol 11, 849–60 (2010).

2. Venkei, Z.G. & Yamashita, Y.M. Emerging mechanisms of asymmetric stem cell division. J Cell Biol 217, 3785–3795 (2018).

3. Clevers, H. Stem cells, asymmetric division and cancer. Nat Genet 37, 1027–8 (2005).

4. Morrison, S.J. & Kimble, J. Asymmetric and symmetric stem-cell divisions in development and cancer. Nature 441, 1068–74 (2006).

5. Kahney, E.W., Ranjan, R., Gleason, R.J. & Chen, X. Symmetry from Asymmetry or Asymmetry from Symmetry? Cold Spring Harb Symp Quant Biol 82, 305–318 (2017).

6. Tran, V., Lim, C., Xie, J. & Chen, X. Asymmetric division of Drosophila male germline stem cell shows asymmetric histone distribution. Science 338, 679–82 (2012).

7. Betschinger, J. & Knoblich, J.A. Dare to be different: asymmetric cell division in Drosophila, C. elegans and vertebrates. Curr Biol 14, R674–85 (2004).

8. Inaba, M. & Yamashita, Y.M. Asymmetric stem cell division: precision for robustness. Cell Stem Cell 11, 461–9 (2012).

9. Tarayrah, L. & Chen, X. Epigenetic regulation in adult stem cells and cancers. Cell Biosci 3, 41 (2013).

10. Wu, H. & Sun, Y.E. Epigenetic regulation of stem cell differentiation. Pediatr Res 59, 21R–5R (2006).

11. Wutz, A. Epigenetic regulation of stem cells: the role of chromatin in cell differentiation. Adv Exp Med Biol 786, 307–28 (2013).

12. Spradling, A., Fuller, M.T., Braun, R.E. & Yoshida, S. Germline stem cells. Cold Spring Harb Perspect Biol 3, a002642 (2011).

13. Fuller, M.T. & Spradling, A.C. Male and female Drosophila germline stem cells: two versions of immortality. Science 316, 402–4 (2007).

14. Tran, V., Feng, L. & Chen, X. Asymmetric distribution of histones during Drosophila male germline stem cell asymmetric divisions. Chromosome Res 21, 255–69 (2013).

15. Alabert, C. & Groth, A. Chromatin replication and epigenome maintenance. Nat Rev Mol Cell Biol 13, 153–67 (2012).

16. Bellush, J.M. & Whitehouse, I. DNA replication through a chromatin environment Philos Trans R Soc Lond B Biol Sci 372 (2017).

17. Gunesdogan, U., Jackle, H. & Herzig, A. Histone supply regulates S phase timing and cell cycle progression. Elife 3, e02443 (2014).

18. McKnight, S.L. & Miller, O.L., Jr. Electron microscopic analysis of chromatin replication in the cellular blastoderm Drosophila melanogaster embryo. Cell 12, 795–804 (1977).

19. Sogo, J.M., Stahl, H., Koller, T. & Knippers, R. Structure of replicating simian virus 40 minichromosomes. The replication fork, core histone segregation and terminal structures. J Mol Biol 189, 189–204 (1986).

20. Ramachandran, S. & Henikoff, S. Replicating Nucleosomes. Sci Adv 1(2015).

21. Burgess, R.J. & Zhang, Z. Histone chaperones in nucleosome assembly and human disease. Nat Struct Mol Biol 20, 14–22 (2013).

22. Serra-Cardona, A. & Zhang, Z. Replication-Coupled Nucleosome Assembly in the Passage of Epigenetic Information and Cell Identity. Trends Biochem Sci 43, 136–148 (2018).

23. Grover, P., Asa, J.S. & Campos, E.I. H3-H4 Histone Chaperone Pathways. Annu Rev Genet 52, 109–130 (2018).

24. Seale, R.L. Studies on the mode of segregation of histone nu bodies during replication in HeLa cells. Cell 9, 423–9 (1976).

25. Leffak, I.M., Grainger, R. & Weintraub, H. Conservative assembly and segregation of nucleosomal histones. Cell 12, 837–45 (1977).

26. Riley, D. & Weintraub, H. Conservative segregation of parental histones during replication in the presence of cycloheximide. Proc Natl Acad Sci U S A 76, 328–32 (1979).

27. Seidman, M.M., Levine, A.J. & Weintraub, H. The asymmetric segregation of parental nucleosomes during chrosome replication. Cell 18, 439–49 (1979).

28. Weintraub, H. Cooperative alignment of nu bodies during chromosome replication in the presence of cycloheximide. Cell 9, 419–22 (1976).

29. Petryk, N. et al. MCM2 promotes symmetric inheritance of modified histones during DNA replication. Science 361, 1389–1392 (2018).

30. Yu, C. et al. A mechanism for preventing asymmetric histone segregation onto replicating DNA strands. Science (2018).

31. Roufa, D.J. & Marchionni, M.A. Nucleosome segregation at a defined mammalian chromosomal site. Proc Natl Acad Sci U S A 79, 1810–4 (1982).

32. Snedeker, J., Wooten, M. & Chen, X. The Inherent Asymmetry of DNA Replication. Annu Rev Cell Dev Biol 33, 291–318 (2017).

33. Sheng, X.R. & Matunis, E. Live imaging of the Drosophila spermatogonial stem cell niche reveals novel mechanisms regulating germline stem cell output. Development 138, 3367–76 (2011).

34. Yadlapalli, S., Cheng, J. & Yamashita, Y.M. Drosophila male germline stem cells do not asymmetrically segregate chromosome strands. J Cell Sci 124, 933–9 (2011).

35. Yadlapalli, S. & Yamashita, Y.M. Chromosome-specific nonrandom sister chromatid segregation during stem-cell division. Nature (2013).

36. Xie, J. et al. Histone H3 Threonine Phosphorylation Regulates Asymmetric Histone Inheritance in the Drosophila Male Germline. Cell 163, 920–33 (2015).

37. Cohen, S.M., Chastain, P.D., 2nd, Cordeiro-Stone, M. & Kaufman, D.G. DNA replication and the GINS complex: localization on extended chromatin fibers. Epigenetics Chromatin 2, 6 (2009).

38. Ahmad, K. & Henikoff, S. Histone H3 variants specify modes of chromatin assembly. Proc Natl Acad Sci U S A 99 Suppl 4, 16477–84 (2002).

39. Blower, M.D., Sullivan, B.A. & Karpen, G.H. Conserved Organization of Centromeric Chromatin in Flies and Humans. Developmental Cell 2, 319–330 (2002).

40. Wolstenholme, D.R. Replicating DNA molecules from eggs of Drosophila melanogaster. Chromosoma 43, 1–18 (1973).

41. Blumenthal, A.B., Kriegstein, H.J. & Hogness, D.S. The units of DNA replication in Drosophila melanogaster chromosomes. Cold Spring Harb Symp Quant Biol 38, 205–23 (1974).

42. Ke, M.T. et al. Super-Resolution Mapping of Neuronal Circuitry With an Index-Optimized Clearing Agent. Cell Rep 14, 2718–32 (2016).

43. Hell, S.W. & Wichmann, J. Breaking the diffraction resolution limit by stimulated emission: stimulated-emission-depletion fluorescence microscopy. Opt Lett 19, 780–2 (1994).

44. Blythe, S.A. & Wieschaus, E.F. Zygotic genome activation triggers the DNA replication checkpoint at the midblastula transition. Cell 160, 1169–81 (2015).

45. Wold, M.S. Replication protein A: a heterotrimeric, single-stranded DNA-binding protein required for eukaryotic DNA metabolism. Annu Rev Biochem 66, 61–92 (1997).

46. Alabert, C. et al. Two distinct modes for propagation of histone PTMs across the cell cycle. Genes Dev 29, 585–90 (2015).

47. Petryk, N. et al. MCM2 promotes symmetric inheritance of modified histones during DNA replication. Science (2018).

48. Horvath, P. & Barrangou, R. CRISPR/Cas, the immune system of bacteria and archaea. Science 327, 167–70 (2010).

49. Wright, A.V., Nunez, J.K. & Doudna, J.A. Biology and Applications of CRISPR Systems: Harnessing Nature’s Toolbox for Genome Engineering. Cell 164, 29–44 (2016).

50. Lasko, P.F. & Ashburner, M. Posterior localization of vasa protein correlates with, but is not sufficient for, pole cell development. Genes Dev 4, 905–21 (1990).

51. Annunziato, A.T. Assembling chromatin: the long and winding road. Biochim Biophys Acta 1819, 196–210 (2013).

52. Jackson, V. & Chalkley, R. A new method for the isolation of replicative chromatin: selective deposition of histone on both new and old DNA. Cell 23, 121–34 (1981).

53. Jackson, V. & Chalkley, R. Histone segregation on replicating chromatin. Biochemistry 6930–8 (1985).

54. Martin-Parras, L., Hernandez, P., Martinez-Robles, M.L. & Schvartzman, J.B. Unidirectional replication as visualized by two-dimensional agarose gel electrophoresis. J Mol Biol 220, 843–53 (1991).

55. Dalgaard, J.Z. & Klar, A.J. A DNA replication-arrest site RTS1 regulates imprinting by determining the direction of replication at mat1 in S. pombe. Genes Dev 15, 2060–8 (2001).

56. Sasaki, T., Sawado, T., Yamaguchi, M. & Shinomiya, T. Specification of regions of DNA replication initiation during embryogenesis in the 65-kilobase DNApolalpha-dE2F locus of Drosophila melanogaster. Mol Cell Biol 19, 547–55 (1999).

57. Buck, S.W., Sandmeier, J.J. & Smith, J.S. RNA polymerase I propagates unidirectional spreading of rDNA silent chromatin. Cell 111, 1003–14 (2002).

58. Hand, R. Regulation of DNA replication on subchromosomal units of mammalian cells. J Cell Biol 64, 89–97 (1975).

59. Huberman, J.A. & Tsai, A. Direction of DNA replication in mammalian cells. J Mol Biol 75. 5–12 (1973).

60. Palmigiano, A. et al. PREP1 tumor suppressor protects the late-replicating DNA by controlling its replication timing and symmetry. Sci Rep 8, 3198 (2018).

