## Supplementary for "Unidirectional fork movement coupled with strand-specific histone incorporation ensures asymmetric histone inheritance"

### Supplementary Materials

### Materials and Methods

#### Fly strains and husbandry

Fly stocks were raised using standard Bloomington medium at 18°C, 25°C, or 29°C as noted. The following fly stocks were used: *hs-flp* on the X chromosome (Bloomington Stock Center BL-26902), *nos-Gal4* on the 2nd chromosome[1], *UASp-FRT-H3-GFP-PolyA-H3-mKO* on the 3rd chromosome and *UASp-FRT-H2B-GFP-PolyA-H2B-mKO,* as reported previously[2]. Other new histone transgenic strains generated for this work are described as follows and are all on either the 2nd or the 3rd chromosome as a single-copy transgene.

#### Generation of fly strains with different switchable dual-color transgenes

Standard procedures were used for all molecular cloning experiments. Enzymes used for plasmid construction were obtained from New England Biolabs (Beverly, MA). The new histone sequences, including *histone-mKO, histone-mCherry* or *histone-GFP*, were recovered as an XbaI flanked fragment and were subsequently inserted into the XbaI site of the UASp plasmid to construct the *UASp-new histone* plasmid. The old histone sequences, including *histone-GFP, histone-EGFP*, or *histone-mKO,* were inserted to *pBluescript-FRT-NheI-SV40 PolyA-FRT* plasmid at the unique NheI site. The entire *NotI-FRT-old histone-SV40 PolyA-FRT-EcoRI* sequences were then subcloned into the *UASp-new histone* plasmid digested by *NotI* and *EcoRI*. The final *UASp-FRT-old histone-PolyA-FRT-new histone* plasmids were introduced to $w^{1118}$ flies by P-element-mediated germline transformation (Bestgene Inc.). Transgenic flies with the following transgenes were newly generated in studies reported here:

*UASp-FRT-H4-GFP-PolyA-FRT-H4-mKO*, *UASp-FRT-H2A-GFP-PolyA-FRT-H2A-mKO*, *UASp-FRT-H2A-EGFP-PolyA-FRT-H2A-mCherry*, *UASp-FRT-H1-GFP-PolyA-FRT-H1-mKO*, *UASp-FRT-H3-mKO-PolyA-FRT-H3-GFP*, and *UASp-FRT-H3-EGFP-PolyA-FRT-H3-mCherry*.

To assess the impact of transgene expression on cell cycle dynamics, live cell imaging was performed on fly strains expression transgenic histones (*nos-Gal4; UASp-histone transgene*) *versus* a control fly strain (*nos-Gal4; UASp-tubulin-GFP*). The average GSC cell cycle lengths between these two groups were not significantly different ($P$ = 0.88; students $t$-test): Average time= 850.0 minutes per GSC cell cycle for the fly lines with histone transgenes (from M-phase of one cell cycle to the subsequent M-phase; $n$ = 10) *versus* Average time= 842.5minutes per GSC cell cycle for the control (from M-phase of one cell cycle to the subsequent M-phase; $n$ = 12).

**Generating knock-in fly strains to tag genes encoding key DNA replication components**

In collaboration with Fungene Inc. (Beijing, China), the following fly line was generated using the CRISPR-Cas9 technology: CG5602 (DNA ligase I, major replicative ligase) with 3xHA tag at the 3' immediately upstream of the STOP codon, generating the fusion protein: DNA ligase-3HA.

**Heat shock scheme**

Flies with *UASp*-dual color histone transgenes were paired with *nos-Gal4* drivers. Flies were raised mostly at 18°C throughout development until adulthood to avoid pre-flipping. In all experiments, flies without heat shock were always checked to evaluate pre-flip events and samples showing pre-flipping activity were excluded from all data acquisition.

For adult males: Before heat shock, 0-3 day old males were transferred to vials that had been air dried for 24 hours. Vials were submerged underneath water up to the plug in a circulating 37°C water bath for 90 minutes and recovered in a 29°C incubator for indicated time before dissection, followed by immunostaining or live cell imaging experiments.

For wandering third-instar larvae: bottles containing third instar larvae (pre-wandering stage) were submerged underneath water up to the plug in a circulating 37°C water bath for 90 minutes and recovered in a 29°C incubator for indicated time before dissection, followed by fiber preparation and immunostaining experiments.

**Immunostaining experiments**

Immunofluorescence staining was performed using standard procedures[2,3]. Primary antibodies used were mouse anti-Fas III (1:200, DSHB, 7G10), anti-HA (1:200; Sigma-Aldrich H3663), anti-PCNA (1:200; Santa Cruz sc-56), anti-GFP (1:1,000; Abcam ab 13970), anti-mKO (1:200; MBL PM051M), anti-mCherry (1:1000; Invitrogen M11217), anti-H3K27me3 (1:200; Millipore 07-449), anti-H4K20me2/3 (1:400; Abcam ab7817), anti-ssDNA (1:100, DSHB) and anti-BrdU (1:200; Abcam ab6326). BrdU analogue was Invitrogen B23151 5-bromo-2´-deoxyuridine (BrdU). Secondary antibodies were the Alexa Fluor-conjugated series (1:1000; Molecular Probes). Confocal images for immunostained fixed sample were taken using Zeiss LSM 700 Multiphoton confocal microscope with 63x or 100x oil immersion objectives and processed using Adobe Photoshop software.

**Quantification of GFP and mKO intensity in whole testis**

No antibody was added to enhance either GFP or mKO signal. Values of GFP and mKO intensity were calculated using Image J software: DAPI signal was used to determine the area of nucleus for measuring both GFP and mKO fluorescent signals, the raw reading was subsequently adjusted by subtracting fluorescence signals in the hub region used as background in both GSC and GB nuclei and compared between each other.

**Chromatin fiber preparation with nucleoside analogue incorporation and immunostaining**

Testes were dissected in Schneider's *Drosophila* medium (Gibco™, Catalog# 21720001) at room temperature (RT) and incubated in Schneider's medium containing 100μM EdU analogue (Invitrogen™ Click-iT™ EdU Imaging Kit, Catalog# C10340). Testes were incubated for 30 minutes, rotating, at RT unless otherwise specified in the protocol. At the end of the 30 minutes, testes were washed for three times with Schneider's medium at RT. Following wash, testes were incubated in the dissociation buffer [Dulbecco's PBS with $Mg^{2+}$ and $Ca^{2+}$ with collagenase/dispase (MilliporeSigma®) added to a final concentration of 2mg/ml] in a 37°C water bath for five minutes. Cells were pelleted at 1000G for five minutes, after which the dissociation buffer was drained. Cells were suspended in 60μl of lysis buffer (80mM NaCl, 150mM Tris-base, 0.2% Joy detergent, pH 10). Following resuspension, 20μl of lysis buffer/cell mixture was transferred to a clean glass slide (Fisherbrand™ Superfrost™ Plus Microscope Slides) and allowed to sit in lysis buffer until cells were fully lysed (~5 minutes). 10μl of sucrose/formalin solution (1M sucrose; 10% formaldehyde) was then added and incubated for two minutes. A coverslip (Fisherbrand™ Microscope Cover Glass, 12-545-J 24x60mm) was placed on top of the lysed chromatin solution, after which, the slide was transferred immediately to liquid nitrogen and allowed to sit there for two minutes. Cover slip was then removed with a razor blade, after which the slide was transferred to cold (-20°C) 95% ethanol for ten minutes. Next, slide was incubated with fixative solution [0.5% formaldehyde in 1xPBST (1xPBS with

0.1% Triton)] for one minute. The fixative solution was drained and the slides were placed into a coplin jar containing 50ml 1xPBS. Slides were washed twice with 50ml 1xPBS each time and placed in a humid chamber with 1ml of blocking solution (2% BSA in 1xPBST) for 30-minute pre-blocking. Blocking buffer was then drained and primary antibodies were added for incubation overnight at 4°C. Slides were then washed twice with 50ml 1xPBS each time and incubated with secondary antibodies for two hours at RT. Slides were then washed twice with 50ml 1xPBS each time and mounted with ProLong® Diamond mounting media.

For BrdU labeling, fibers were treated with 1M HCL for two hours at 37°C to expose BrdU epitope prior to addition of anti-BrdU primary antibody. For EdU visualization: EdU analogue was conjugated to Alexa-647 dye using CLICK chemistry (reviewed by [4,5]).

**Identification of replicating regions at chromatin fibers**

Replicating regions were identified by EdU incorporation into the chromatin fiber. Replication bubbles were identified as EdU-positive regions showing a transition from a single fiber structure to a double fiber structure, which was co-localized with EdU incorporation. EdU-positive regions along the chromatin fiber were also associated with a significant increase in DNA dye intensity, referred to as DAPI-bright regions. DAPI-bright regions were identified as regions showing greater than a 2-fold increase in DAPI-intensity relative to surrounding DAPI-dim regions from the same chromatin fiber. This greater than 2-fold difference in DAPI intensity likely reflects both the two-fold increase in DNA content associated with replication and the differences in super-helical torsion associated with the advancing replication fork. Previous studies have reported that negative-supercoiling is enriched behind the replication fork, whereas positive-supercoiling is enriched ahead of the replication fork[6-8]. The unwound nature of the negatively supercoiled DNA favors the binding of intercalating DNA dyes, such as DAPI and Yoyo-III[9,10]. Conversely, positive supercoiling structure limits the accessibility of DNA to intercalating small molecules.

**Superresolution STED imaging**

Superresolution images were acquired using a Leica TCS SP8 STED microscope with a 1.4 NA 100X STED white objective. Immunostaining experiments were performed to enhance specimen brightness and photo-stability for STED microscopy. Secondary antibody fluorophore conjugates were empirically selected for STED performance. The optimal 3-colour separation was performed with the 592 nm continuous wave (CW) and 775 nm pulsed depletion lasers (Alexa 488 with STED 592nm, Alexa 568 with STED 775nm, and Alexa 647 with STED 775nm). Images were acquired as single z-planes for all tissue types, including whole mount and squash tissues, as well as isolated fibers, in order to minimize drift between channel acquisitions. Specimens included 100 nm TetraSpeck microsphere beads as fiducial markers (Thermo Fisher Catalog# T7279). Instrument aberration and blurring was corrected with post-acquisition deconvolution using the Scientific Volume Imagine (SVI) Huygens Professional software package, which achieved improved calculated/theoretical PSFs *via* complete integration with the Leica LAS-X software and hardware. Detailed instrument acquisition and post-processing settings are available upon request.

**Quantification of proteins on sister chromatids without strandness information**

All fiber analyses were performed using Java image processing program FIJI. To capture localized distribution of histones and other proteins on chromatin fibers, images were imported into FIJI and line plots were drawn across sister chromatids to measure average fluorescence intensity at the specified region. To measure histone distribution differences between sister chromatids, replication regions longer than 2μm in length were subdivided into 2μm-long segments along the length of the chromatin fiber. Two microns were chosen as this was the average size of individual replicons with 30 minute EdU pulses (Supplementary figure 3). Given the estimated average rate of DNA polymerase to synthesize ~0.5- 2.0 kb DNA per minute[11], this 2μm chromatin fiber reflects approximately 15-60 kb of genomic DNA. For replication region shorter than 2μm, the entire length of the region containing resolvable sister chromatids was used to asses differences in histone distribution. To effectively compare histone distribution patterns across multiple data sets, we normalized them using the following strategy: First, we quantified fluorescence levels for histone signals [e.g. old histones (GFP or EGFP), new histones (mKO or mCherry)] for each sister chromatid fiber segment. We then divided fluorescence intensity from the brighter sister chromatid fiber segment by the fluorescence intensity from the less bright sister chromatid fiber segment, to generate a ratio of the relative difference between sister chromatids. This quantification scheme was used for Figure 2 and Supplemental figure 5.

**Classification of different categories of chromatin fibers without strandness information**

For old histone on sister chromatids, we classified fibers as symmetric (ratio <1.80), moderately asymmetric (1.8< ratio <2.44), or highly asymmetric (ratio >2.44). We used 2.44 as a standard for calling highly asymmetric, as it is two standard deviations above the average ratio observed for old H2A between sister chromatids. For new histones on sister chromatids, we classified fibers as symmetric (ratio <1.70), moderate asymmetric class (1.70< ratio<2.16), or highly asymmetric (ratio >2.16). We used 2.16 as a standard for calling highly asymmetric, as it is two standard deviations above the average ratio observed for new H2A between sister chromatids.

**Quantification of proteins on sister chromatids with strandness information**

To capture localized distribution of post-translational histone modifications and other proteins on replicating or newly replicated chromatin fibers, replication region longer than 2μm in length were divided into 2μm-segments, as described above. For replication region shorter than 2μm, the entire resolvable sister chromatids region was used to asses differences in the distribution of post-translational histone modifications. Lagging strand-enriched proteins, such as RPA and PCNA, were used as a proxy for lagging strands. To compare distribution of post-translational histone modifications on replicating or newly replicated chromatin fibers, the leading strand (RPA-depleted or PCNA-depleted strand) was divided by the lagging strand (RPA-enriched or PCNA-enriched) to generate: Ratio= leading strand protein levels/lagging strand protein levels. Ratios were then $\log_2$-transformed such that leading strand enrichment would appear as a positive value, and lagging strand enrichment would appear as a negative value.

To retrieve the information of average fold-enrichment: Average fold enrichment = 2^ (Average of $\log_2$ values).

**Classification of chromatin fibers with strandness information**

To allow for comparison between different data sets of chromatin fibers with leading/lagging strand information, the following criteria were used: Fibers with a $\log_2$ (leading strand/lagging strand ratio)$\geq$ 0.5 were classified as leading strand enriched. Fibers with a -0.5< $\log_2$ (leading strand/lagging strand ratio)< 0.5 were classified as symmetric. Fibers with $\log_2$ (leading strand/lagging strand ratio)$\leq$ -0.5 were classified as lagging strand enriched.

**Sequential incorporation of EdU and BrdU**

EdU labeling was performed using Click-iT Plus EdU Alexa Fluor 647 Imaging Kit (Life Science C10640) according to manufacturer's instructions. Dissected testes were immediately incubated in in Schneider's *Drosophila* medium (Gibco™, Catalog# 21720001) with 100 μM EdU for 30 minutes at RT. The testes were subsequently fixed and incubated with primary antibody, as described above. Fluorophore conjugation to EdU was performed along manufacturer's instructions and followed by secondary antibody incubation.

For double-labelling (EdU followed by BrdU), testes were first incubated for ten minutes in Schneider's medium containing 100μM of EdU. Samples were then washed three times quickly: Washes entailed re-suspending testes in fresh Schneider's medium, allowing testes to settle to the bottom of the eppendorf tube, and then quickly pipetting away extra Schneider's medium. All three washes were completed within two minutes. Testes were then transferred to Schneider's medium containing 100μM of BrdU analog and incubated for ten minutes at RT, after which, testes were rigorously washed three times as described above. Chromatin fibers were then generated as described above and DNA fibers were then generated as following.

**DNA fiber preparation**

Testes were dissected in fresh Schneider's *Drosophila* medium (Gibco™, Catalog# 21720001) at RT and incubated in Schneider’s medium containing 100µM EdU analogue (Invitrogen™ Click-iT™ EdU Imaging Kit, Catalog# C10340). Testes were incubated for 30 minutes, rotating, at RT unless otherwise specified in the protocol. At the end of the 30 minutes, testes were washed for three times with Schneider's medium at RT. Following wash, testes were incubated in the dissociation buffer [Dulbecco’s PBS with $Mg^{2+}$ and $Ca^{2+}$ with collagenase/dispase (MilliporeSigma®) added to a final concentration of 2mg/ml] in a 37°C water bath for five minutes. Cells were pelleted at 1000G for five minutes, after which the dissociation buffer was drained. Cells were resuspended in 20µl 1xPBS. 10µl of cell suspension was then transferred to a clean glass slide (Fisherbrand™ Superfrost™ Plus Microscope Slides). Slides were left to sit until all moisture had nearly evaporated. Cells were then incubated in 10µls of DNA lysis buffer (200 mM Tris–HCl, pH 7.5, 50 mM EDTA, 0.5% SDS). Slides were then tilted at a 25° angle and allow the drop to slowly travel down the slide. Slides were dried for 10 minutes at RT, after which they were fixed in freshly prepared methanol:acetic acid (v/v 3:1) for five minutes. Slides were allowed to dry again (~ 10 minutes) after which slides were washed with 1xPBS. Slides were then stained for EdU, BrdU, or anti-ssDNA antibody that recognizes all DNA after HCl treatment[12].

**DNA fiber staining**

Slides were washed with 1xPBS and then transferred to coplin jar containing 50ml of 1M HCL. Slides were incubated for 20hrs at 37°C to expose BrdU epitope for immunostaining. To minimize cross-reactivity of BrdU antibody to EdU, visualization of EdU was done using Click-iT Plus EdU Alexa Fluor 568 Imaging Kit (Life Science Catalog# C10640) according to manufacturer's instructions. Following click-chemistry reaction, slides were washed twice in 1xPBS and then blocked for 30 minutes with 2.5% bovine serum albumin (BSA). Following blocking, 200 μls of anti-BrdU antibody was added to the slide. Slides were placed in a humid chamber and incubated overnight at 4°C. Slides were washed in 1xPBS and then blocked for 30 minutes in 5% normal goat serum (NGS). Secondary antibody that recognizes anti-BrdU was added and slides were incubated for 2hrs at RT in a humid chamber. Slides were then washed with 1xPBS and blocked for 30 minutes with 5% BSA. Following blocking, 200μls of anti-ssDNA antibody was added to the slide and incubated in a humid chamber for 1 hr at RT. Slides were washed in 1xPBS and blocked for 30 minutes with 5% NGS. Secondary antibody that recognizes anti-ssDNA was then added and slides were incubated at RT for 30 minutes in a humid chamber. Slides were washed in 1xPBS, dried and mounted with 20μls of Vectashield® mounting medium.

**Determining fork movement in chromatin fibers and DNA fibers**

Linearized fibers containing multiple replicons were analyzed to determine fork movement patterns in chromatin fibers and DNA fibers. Bidirectional replicons were identified by the presence of the early label (EdU) flanked by the late label (BrdU) at both sides (e.g. Figure 4c, 4e, 4g). Unidirectional replicons were identified by an alternating pattern of early (EdU) and late (BrdU) along the length of the fiber (e.g. Figure 4d, 4f, 4h). Only fibers with multiple identified replicons were included for data analysis. DNA labels (e.g. DAPI, Yoyo-III or anti-ssDNA) were always included to ensure the continuity of the analyzed fibers.

**PLA assay**

Following incubation with primary antibodies, proximity ligation assay (PLA) was performed using 20 μL of reaction per step per slide according to Sigma-Aldrich Duolink In Situ PLA manufacturer's instruction (Catalog# DUO92101). In brief, two PLA secondary probes, anti-mouse MINUS (e.g. targeting anti-HA mouse primary) and anti-rabbit PLUS (e.g. targeting either anti-GFP or anti-mKO rabbit primaries), were diluted 1:5 in Ab Diluent buffer provided by manufacturer and incubated overnight at 4 °C. Slides were washed in 1X wash buffer A for 10 minutes, followed by the ligation reaction, in which PLA ligation stock was diluted 1:5 in $dH_2O$ with ligase (added at 1:40), followed by incubation for one hour at 37 °C. Slides were then washed in wash buffer A for five minutes, followed by addition of the PLA amplification reaction (1:5 amplification stock and 1:80 polymerase diluted in $dH_2O$) and incubated for two hours at 37 °C. Slides were then washed with 1X wash buffer B for ten minutes, 0.01X wash buffer B for one minute, and 1X PBS for one minute. Following washes, 100μls anti-FasIII (hub cell marker) was then added to slides and allowed to incubate for 30 minutes at RT. Slides were then washed with 1X PBS, after which anti-mouse secondary (Alexa Fluor 405; 1:1000 Molecular Probes) was added to recognize anti-FasIII (labeling hub cells) and anti-PCNA/anti-HA (labeling S phase cells). Slides were incubated for 2hrs at RT and then washed in 1xPBS and mounted. Images were taken using the Zeiss LSM 700 Multiphoton confocal microscope with a 63× oil immersion objectives and processed using Adobe Photoshop software.

**Supplementary Figures and Figure Legends:**

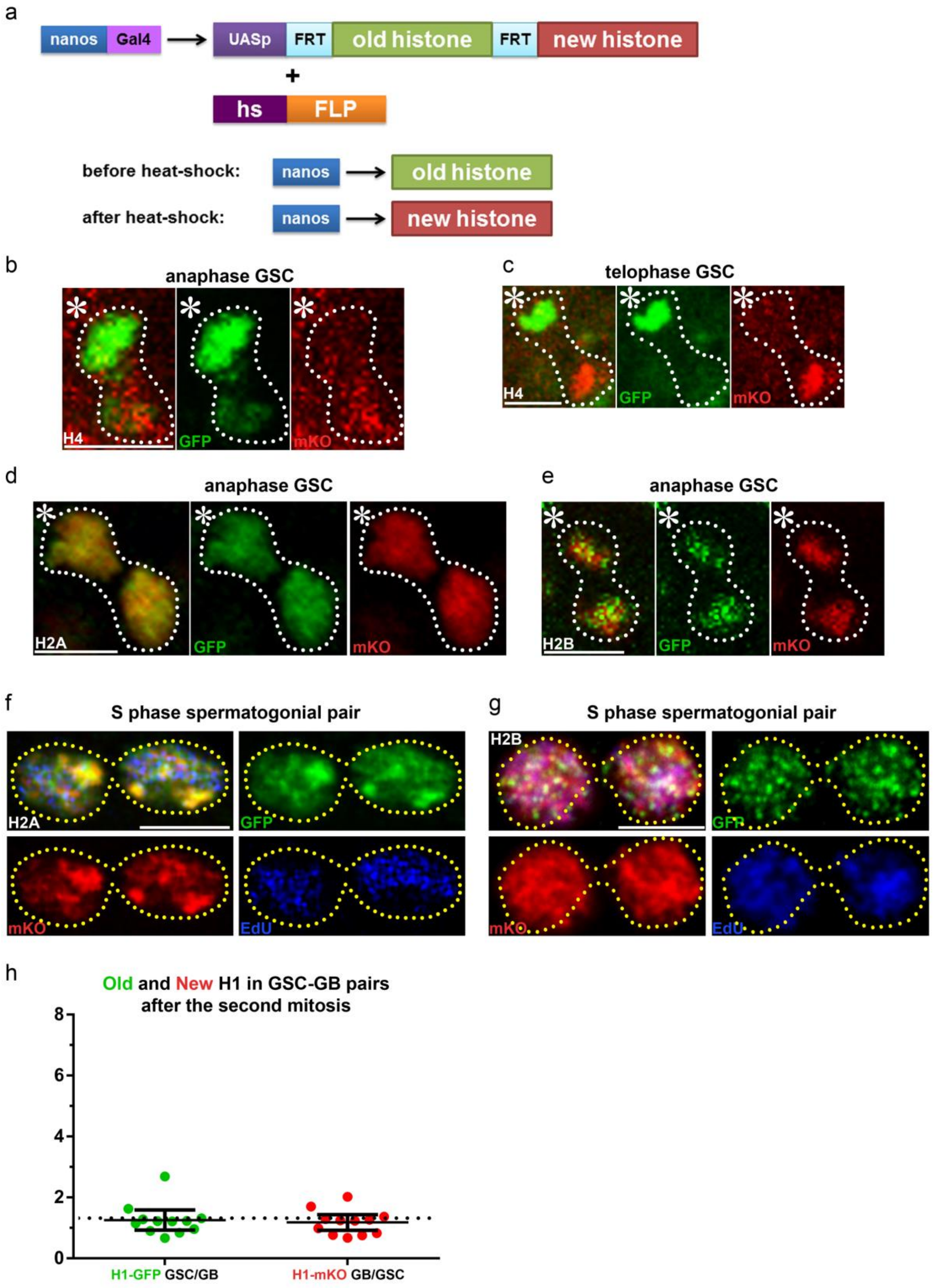

**Supplementary figure 1**: (**a**) A schematic diagram showing the dual color switch design that expresses pre-existing histone and newly synthesized histone by heat-shock treatment, as adapted from[2]. (**b**) An anaphase GSC showing asymmetric segregation of H4-GFP towards the GSC and H4-mKO towards the GB. (**c**) A telophase GSC showing asymmetric distribution of H4-GFP towards the GSC and H4-mKO towards the GB. (**d**) An anaphase GSC showing symmetric segregation of H2A-GFP and H2A-mKO. (**e**) An anaphase GSC showing symmetric segregation of H2B-GFP and H2B-mKO. (**f**) Symmetric H2A inheritance pattern in a post-mitotic SG pair. (**g**) Symmetric H2B inheritance pattern in a post-mitotic SG pair. Scale bars: 5μm. Asterisk: hub. (**h**) Histone H1 showed overall symmetric inheritance pattern in post-mitotic GSC-GB pairs ($n$=12). See Supplementary table 4 for details. Error bars represent 95% confidence interval. Neither old H1 nor new H1 is significantly different from the value of 1; $P = 0.092$ for old H1; $P = 0.151$ for new H1, based on Wilcoxon signed rank test.

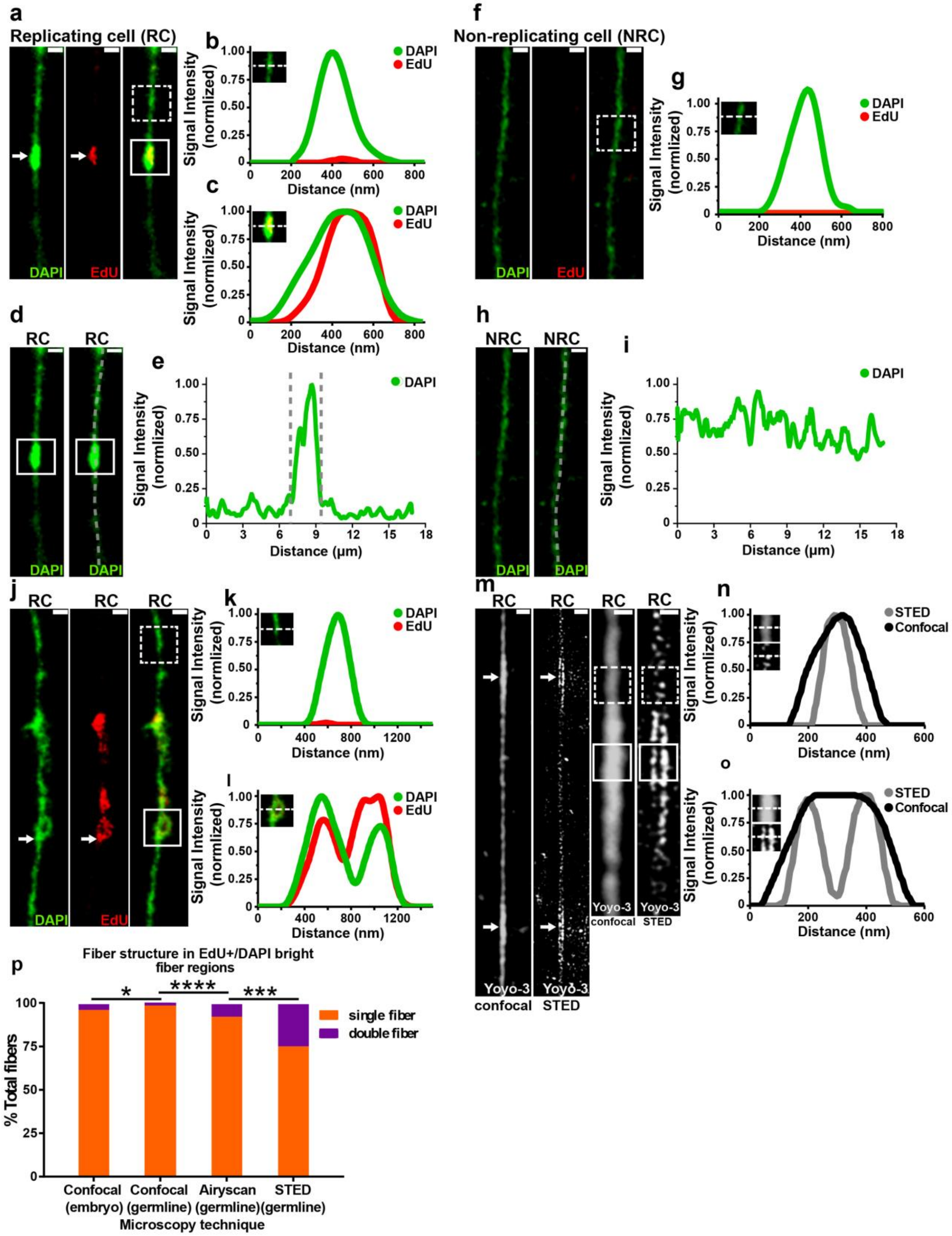

a
Replicating cell (RC)
DAPI
EdU
b
c
Signal Intensity (normlized)
Distance (nm)
DAPI
EdU
d
RC
e
Distance (µm)
f
Non-replicating cell (NRC)
g
h
NRC
i
j
k
l
m
n
o
STED
Confocal
Yoyo-3 confocal
Yoyo-3 STED
Signal Intensity (normalized)
p
Fiber structure in EdU+/DAPI bright fiber regions
% Total fibers
single fiber
double fiber
Confocal (embryo)
Confocal (germline)
Airyscan (germline)
STED (germline)
Microscopy technique

**Supplementary figure 2: Superresolution microscopy helps visualize sister chromatids on isolated chromatin fibers.** (**a**) Confocal image of chromatin fiber isolated from replicating cells (RCs) in the *Drosophila* embryo showing replication "bubble" structure with EdU and brighter DAPI signal (white arrow). (**b**) Line-plot of DAPI and EdU distribution on unreplicated region without EdU (box with dotted white lines in **a**, inset in **b**). (**c**) Line-plot of DAPI and EdU distribution on replicated region with EdU (box with solid white lines in **a**, inset in **c**). Replicated region appears thicker [400nm full-width half maximum (FWHM)] than un-replicated region (250nm FWHM). However, sister chromatids cannot be clearly resolved. (**d**) DAPI-signal from RC-derived chromatin fiber shows brighter DAPI signal in replicating regions (white box). (**e**) Longitudinal line-plot of RC-derived chromatin fiber shows a clear increase in DAPI signal in EdU-positive region, relative to the surrounding EdU-negative region from the same fiber. (**f**) Confocal image of chromatin fiber isolated from non-replicating cells (NRC) in the *Drosophila* adult eye. NRC-derived fibers show uniform DAPI-signal with no detectable EdU staining. (**g**) Line-plot from NRC of DAPI and EdU distribution on randomly selected fiber region (box with dotted white lines in **f**, inset in **g**). (**h**) DAPI shows uniform staining along the length of NRC-derived chromatin fibers. (**i)** Longitudinal line-plot of DAPI intensity in NRC-derived chromatin fiber shows small fluctuations in signal with no significant increases in intensity comparable to those observed in fibers derived from RCs in (**e**). (**j)** Confocal image of chromatin fiber isolated from *Drosophila* embryo shows replication bubble structure. DAPI shows a clear transition from single fiber to double fiber structure at the point where EdU incorporation becomes apparent (white arrow). (**k**) Line-plot of DAPI and EdU distribution on unreplicated region without EdU (box with dotted white lines in **j**, inset in **k**). (**l**) Line plot of DAPI and EdU distribution on replicated region with EdU (box with solid white lines in **j**, inset in **l**). (**m**) Confocal and STED images of chromatin fiber isolated from *Drosophila* male germline, stained with DNA dye Yoyo3. Yoyo bright regions (white arrows) can be resolved into sister chromatids with STED. (**n**) Line plot of Yoyo3 signal in Yoyo3 dim region shows a single fiber structure with both confocal and STED (box with dotted white lines in **m**, inset in **n**). (**o**) Line-plot of Yoyo3 signal in Yoyo3-bright region shows clear double fiber structure with STED but not with confocal (box with solid white lines in **m**, inset in **o**). (**p**) Quantification of the frequency of seeing single *versus* double fiber structure at EdU-positive and DAPI-bright regions. **** $P < 10^{-4}$, *** $P < 10^{-3}$, * $P < 0.05$, Chi-squared test. Scale bars: 500nm.

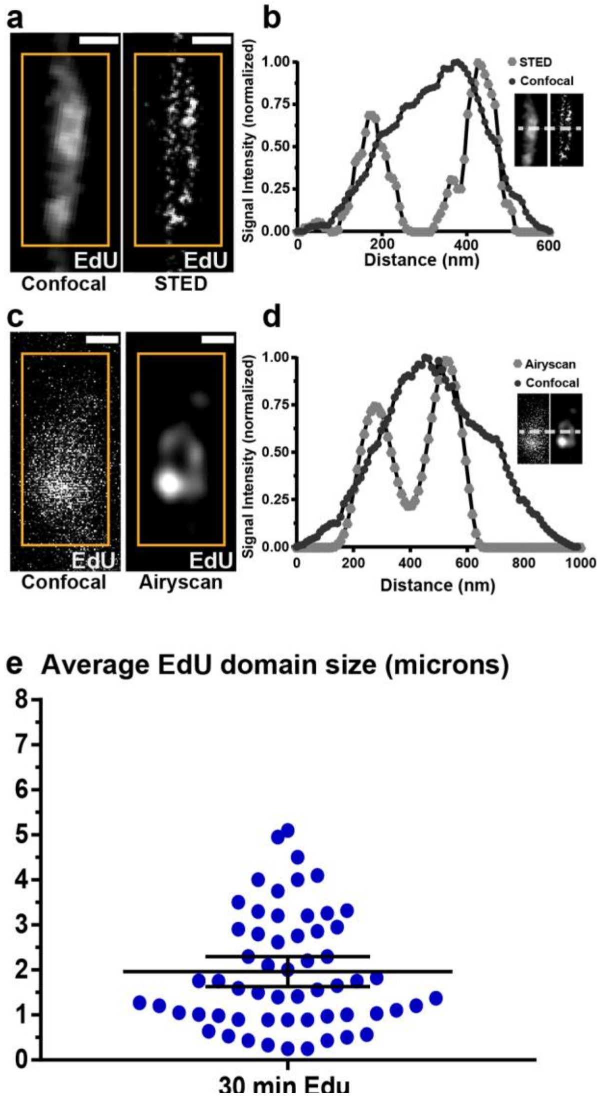

**Supplementary figure 3: Establishment of using chromatin fiber method to quantify differences between sister chromatids.** (**a**) Confocal *versus* STED images of EdU signal on replicating chromatin fiber. The EdU-positive region (box with solid orange lines) cannot be resolved into sister chromatids with confocal but can be resolved with STED. (**b**) Line-plot of EdU signal shows a single fiber structure with confocal imaging but a double fiber structure with STED. (**c**) Confocal *versus* Airyscan images of EdU signal on replicating chromatin fiber. The EdU-positive region (box with solid orange lines) cannot be resolved into sister chromatids with confocal but can be resolved with Airyscan. (**d**) Line-plot of EdU signal shows a single fiber structure with confocal imaging but a double fiber structure with Airyscan. (**e**) Quantification of average EdU-positive regions in replicating chromatin fibers. A 30 minute pulse of EdU incorporation yields an average of 1.96 microns ($n$ = 58) of EdU-positive region. Given the estimated average rate of DNA polymerase to synthesize ~0.5- 2.0 kb DNA per minute[11], this 2μm chromatin fiber reflects approximately 15-60 kb of DNA. Error bars represent 95% confidence interval.

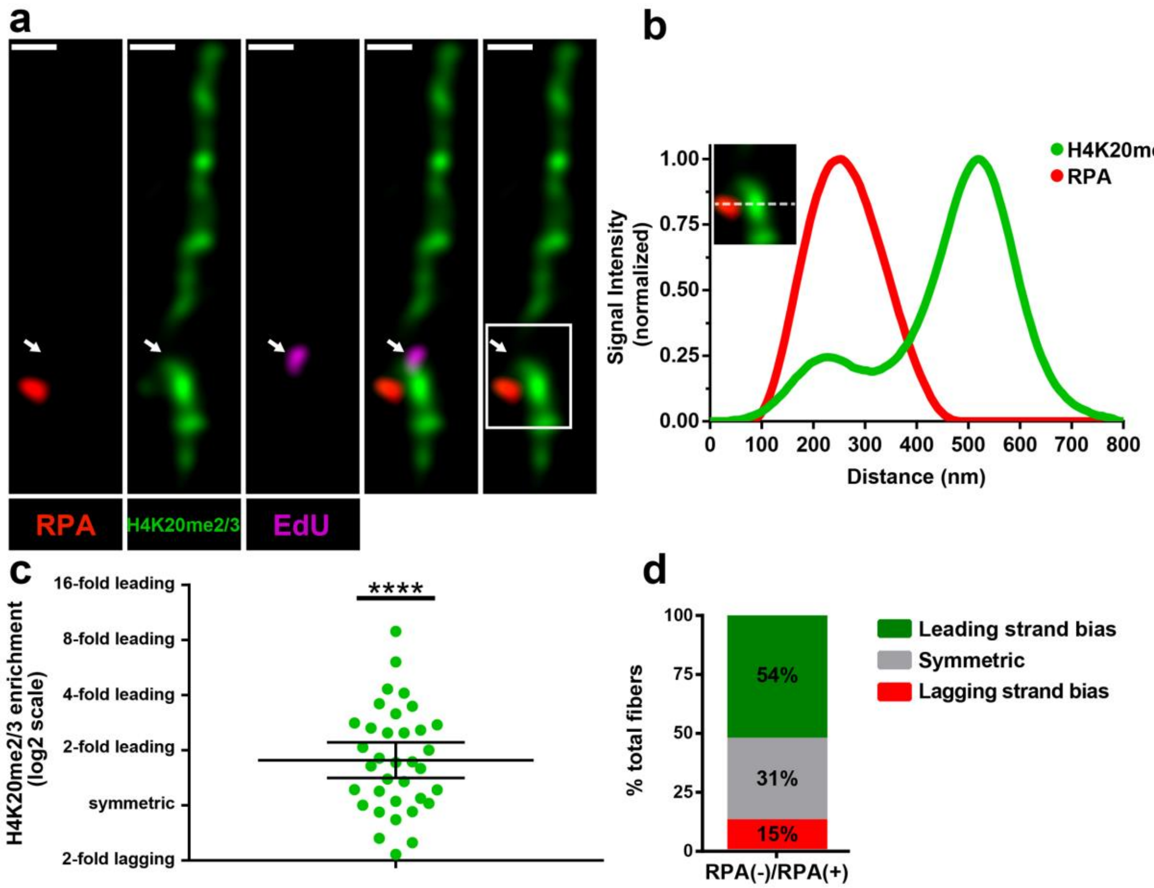

**Supplementary figure 4: Old H4 preferentially associate with the leading strand on chromatin fibers.** (**a**) Airyscan image of a chromatin fiber labeled with EdU (magenta) and H4K20me2/3 (green), RPA (red). The transition from unreplicated single fiber to replicating double fibers is co-localized with the EdU signal (white arrow). H4K20me2/3 signal is enriched at the RPA-depleted sister chromatid. Scale bar: 500nm. (**b**) Line-plot shows H4K20me2/3 and RPA distribution across replicating region (box with solid white lines in **a**, inset in **b**). (**c**) Quantification of the ratio H4K20me2/3 on RPA-depleted sister chromatid/ RPA-enriched sister chromatid (2.06±0.15 *n*=36 replicating regions from 22 chromatin fibers). Data is significantly different from symmetric. Y-axis is with $\log_2$ scale. **** $P < 0.0001$, Mann-Whitney U test. (**d**) Classification of RPA-labeled sister chromatids into 54% leading-strand enriched (ratio >1.4), 15% lagging-strand enriched (ratio <1.4) and 31% symmetric (-1.4< ratio< 1.4).

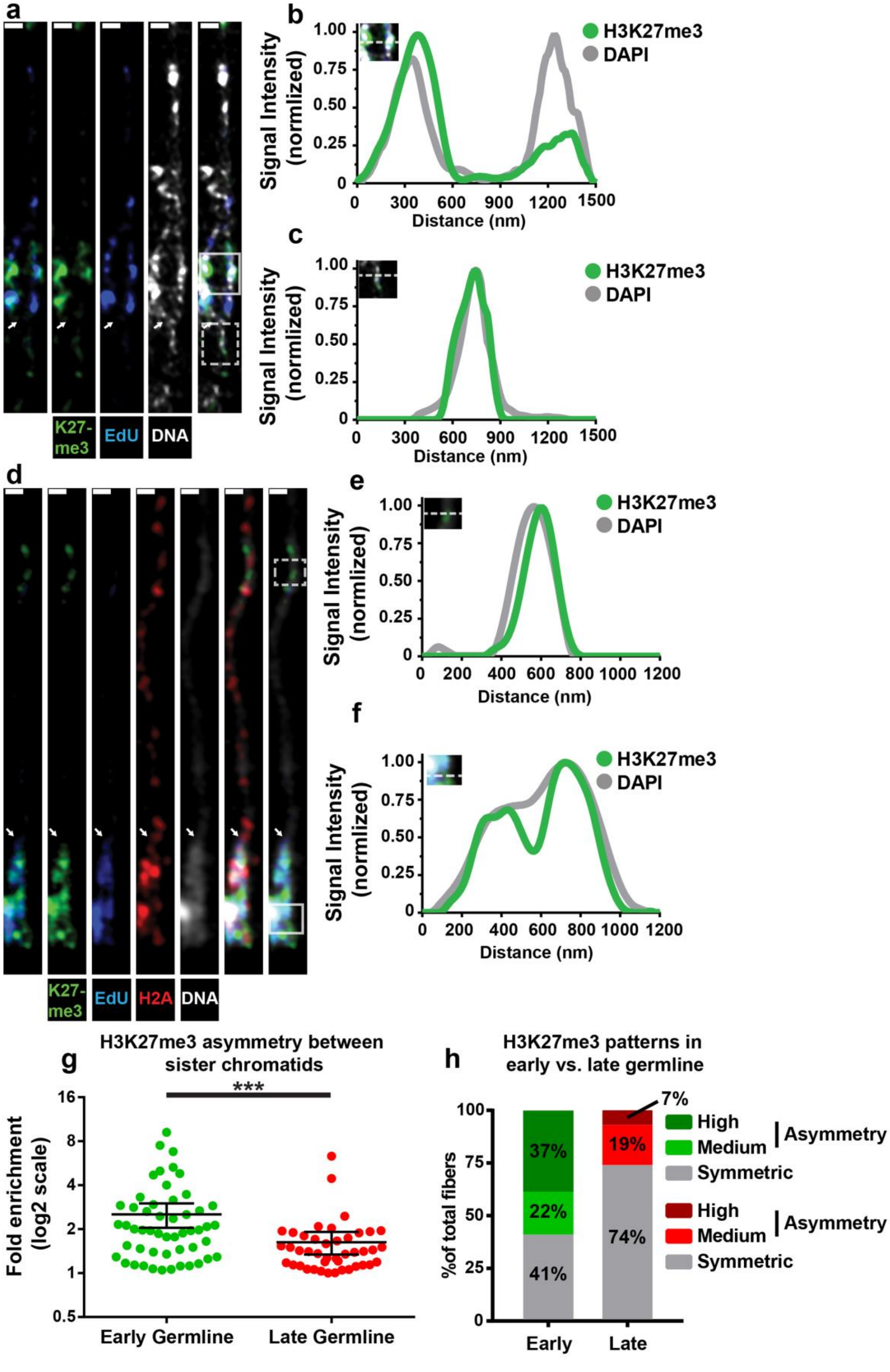
a
K27-me3
EdU
DNA
b
Signal Intensity (normlized)
Distance (nm)
H3K27me3
DAPI
c
Signal Intensity (normlized)
Distance (nm)
H3K27me3
DAPI
d
K27-me3
EdU
H2A
DNA
e
Signal Intensity (normlized)
Distance (nm)
H3K27me3
DAPI
f
Signal Intensity (normlized)
Distance (nm)
H3K27me3
DAPI
g
H3K27me3 asymmetry between sister chromatids
***
Fold enrichment (log2 scale)
Early Germline
Late Germline
h
H3K27me3 patterns in early vs. late germline
%of total fibers
37%
22%
41%
7%
19%
74%
Early
Late
High
Medium
Symmetric
Asymmetry
High
Medium
Symmetric
Asymmetry

**Supplementary figure 5: Comparison of chromatin fibers from early-stage germ cells with those from late-stage germ cells.** (**a**) Airyscan image of a chromatin fiber from early-stage germ cells labeled with EdU (blue) and H3K27me3 (green), DAPI (white). The transition from unreplicated single fiber to replicating double fibers is co-localized with the start of EdU signal (white arrow). H3K27me3 shows a significantly brighter signal on one side of the replication bubble. (**b**) Line-plot shows H3K27me3 and DAPI distribution across replicating region (box with solid white lines in **a**, inset in **b**). (**c**) Line-plot shows H3K27me3 and DAPI distribution across unreplicated region (box with dotted white lines in **a**, inset in **c**). (**d**) Airyscan image of a chromatin fiber from late-stage SGs labeled with EdU (blue) and H3K27me3 (green), DAPI (white). The H2A (red) was specifically expressed using a late-germline-specific *bam-Gal4* driver to label fibers from late-stage germ cells. The transition from unreplicated single fiber to replicating double fibers is co-localized with the EdU signal (white arrow). H3K27me3 signal appears more symmetric between sister chromatids. (**e**) Line-plot shows H3K27me3 and DAPI distribution across unreplicated region (box with dotted white lines in **d**, inset in **e**). (**f**) Line-plot shows H3K27me3 and DAPI distribution across replicating region (box with solid white lines in **d**, inset in **f**). (**g**).Quantification of the H3K27me3 ratio between sister chromatids: H3K27me3 signal in chromatin fibers derived from early-stage germ cells show greater asymmetry between sister chromatids when compared to chromatin fibers derived from late-stage germ cells. On average, early germline fibers showed a ~2.5-fold difference (2.52±0.27, $n$ = 52 replicating regions from 37 chromatin fibers) whereas late germline fibers showed only a 1.6-fold difference (1.60±0.14, $n$ = 43 replicating regions from 23 chromatin fibers). Y-axis is with $\log_2$ scale. **** $P$< 0.0001, Mann-Whitney U test. (**h**) Using the previously mentioned criteria (Figures 2e and 2g, Materials and Methods) to define distinct classes of old histone asymmetries: 39% of early germline fibers showed high asymmetry, 20% showed moderate asymmetry, and 41% showed symmetry. In comparison, only 7% of late germline fibers showed high asymmetry, 19% showed moderate asymmetry and 74% were found to be symmetric. Scale bars: 500nm.

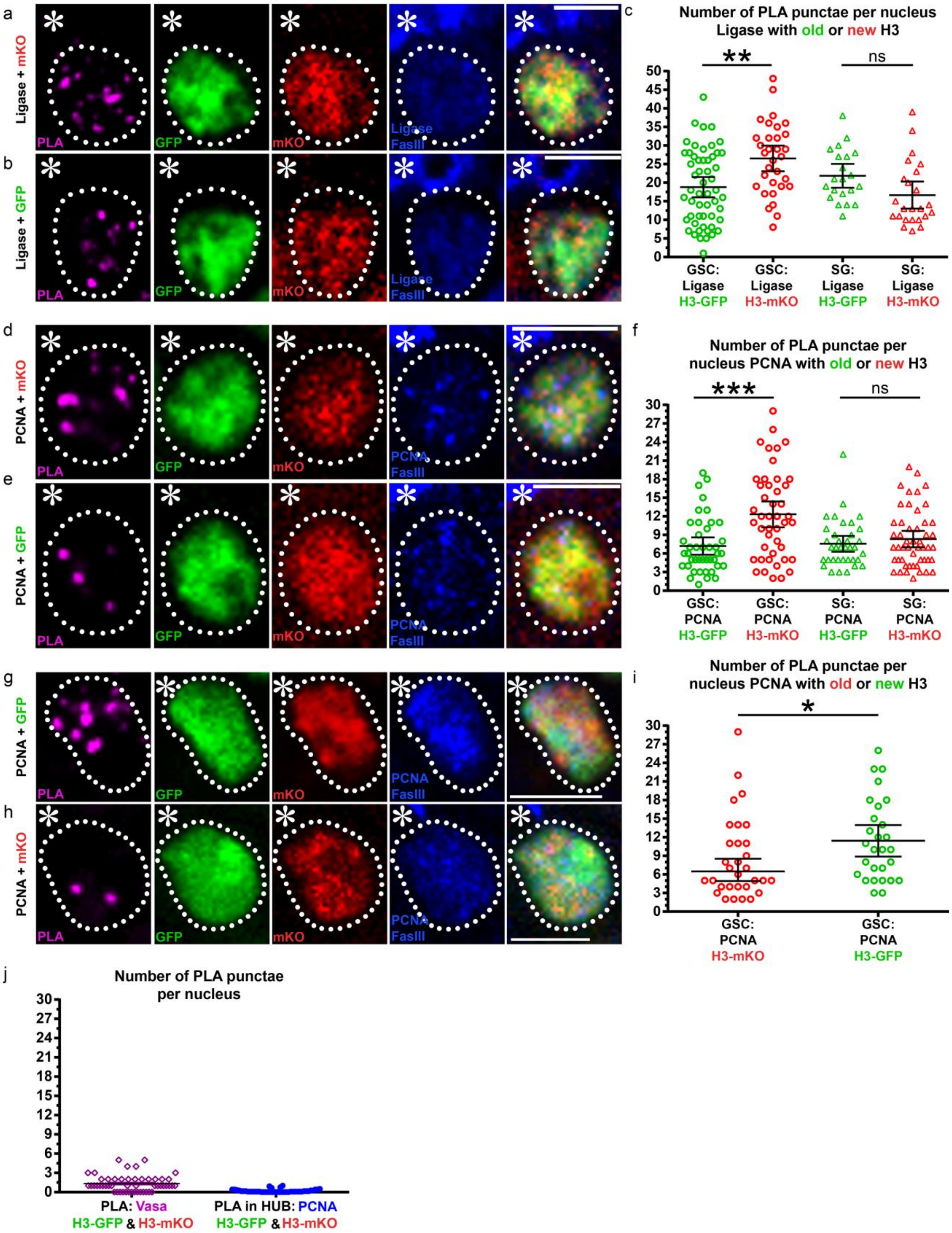
a
Ligase + mKO
PLA
GFP
mKO
Ligase
FasIII
b
Ligase + GFP
c
Number of PLA punctae per nucleus
Ligase with old or new H3
**
ns
GSC: Ligase H3-GFP
GSC: Ligase H3-mKO
SG: Ligase H3-GFP
SG: Ligase H3-mKO
d
PCNA + mKO
PCNA
FasIII
e
PCNA + GFP
f
Number of PLA punctae per nucleus PCNA with old or new H3
***
ns
GSC: PCNA H3-GFP
GSC: PCNA H3-mKO
SG: PCNA H3-GFP
SG: PCNA H3-mKO
g
PCNA + GFP
h
PCNA + mKO
i
Number of PLA punctae per nucleus PCNA with old or new H3
*
GSC: PCNA H3-mKO
GSC: PCNA H3-GFP
j
Number of PLA punctae per nucleus
PLA: Vasa H3-GFP & H3-mKO
PLA in HUB: PCNA H3-GFP & H3-mKO

**Supplementary figure 6: Proximity ligation assay shows distinct proximity between histones (old *versus* new) and lagging strand-enriched DNA replication machinery components in GSCs.** (**a**) A representative GSC showing PLA signals (magenta) between lagging-strand-specific Ligase-HA (blue) and new H3-mKO (red). (**b**) A representative GSC showing PLA signals between Ligase-HA and old H3-GFP (green). (**c**) Quantification of the number of PLA puncta per nucleus between Ligase and histones (old *versus* new) in GSCs and SGs: In GSCs, PLA puntae between ligase and new H3-mKO: 26.5±1.7 (*n*=35); between ligase and old H3-GFP: 18.5±2.5 (*n*=53). In SGs, PLA puntae between ligase and new H3-mKO: 16.7±1.8 (*n*=24); between ligase and old H3-GFP: 21.9±1.5 (*n*=21). (**d**) A representative GSC showing PLA signals (magenta) between lagging-strand enriched PCNA (blue) and new H3-mKO (red). (**e**) A representative GSC showing PLA signals between PCNA and old H3-GFP (green). (**f**) Quantification of the number of PLA puncta per nucleus between PCNA and histones (old *versus* new) in GSCs and SGs: In GSCs, PLA puntae between PCNA and new H3-mKO: 12.3±1.0 (*n*=46); between PCNA and old H3-GFP: 7.2±0.7 (*n*=42). In SGs, PLA puntae between PCNA and new H3-mKO: 10.0±1.8 (*n*=50); between PCNA and old H3-GFP: 7.6±0.6 (*n*=36). **: $P< 0.01$, ***: $P< 0.001$, Kruskal-Wallis multiple comparisons of non-parametric data with Dunn's multiple comparisons corrections test. Error bars represent 95% confidence interval in (**c**) and (**f**). (**g**) A representative GSC showing PLA signals between the lagging strand-enriched component PCNA and new H3-GFP (green). (**h**) A representative GSC showing PLA signals between the lagging strand-enriched component PCNA and old H3-mKO (red). (**i**) Quantification of the number of PLA puncta per nucleus between PCNA and histones (old *versus* new) in GSCs. PLA puntae between PCNA and new H3-GFP: 11.4± 1.2 (*n*=28); between PCNA and old H3-mKO: 8.5±1.2 (*n*=31), *: $P< 0.05$, based on Mann-Whitney U test. (**j)** Quantification of PLA signals in two negative control experiments: first, PLA experiments were performed between histones and a cytoplasmic protein Vasa[13]; second, PLA signals were counted in non-replicating somatic hub cells**.** Both showed very low signals. Scale bars: 5μm.

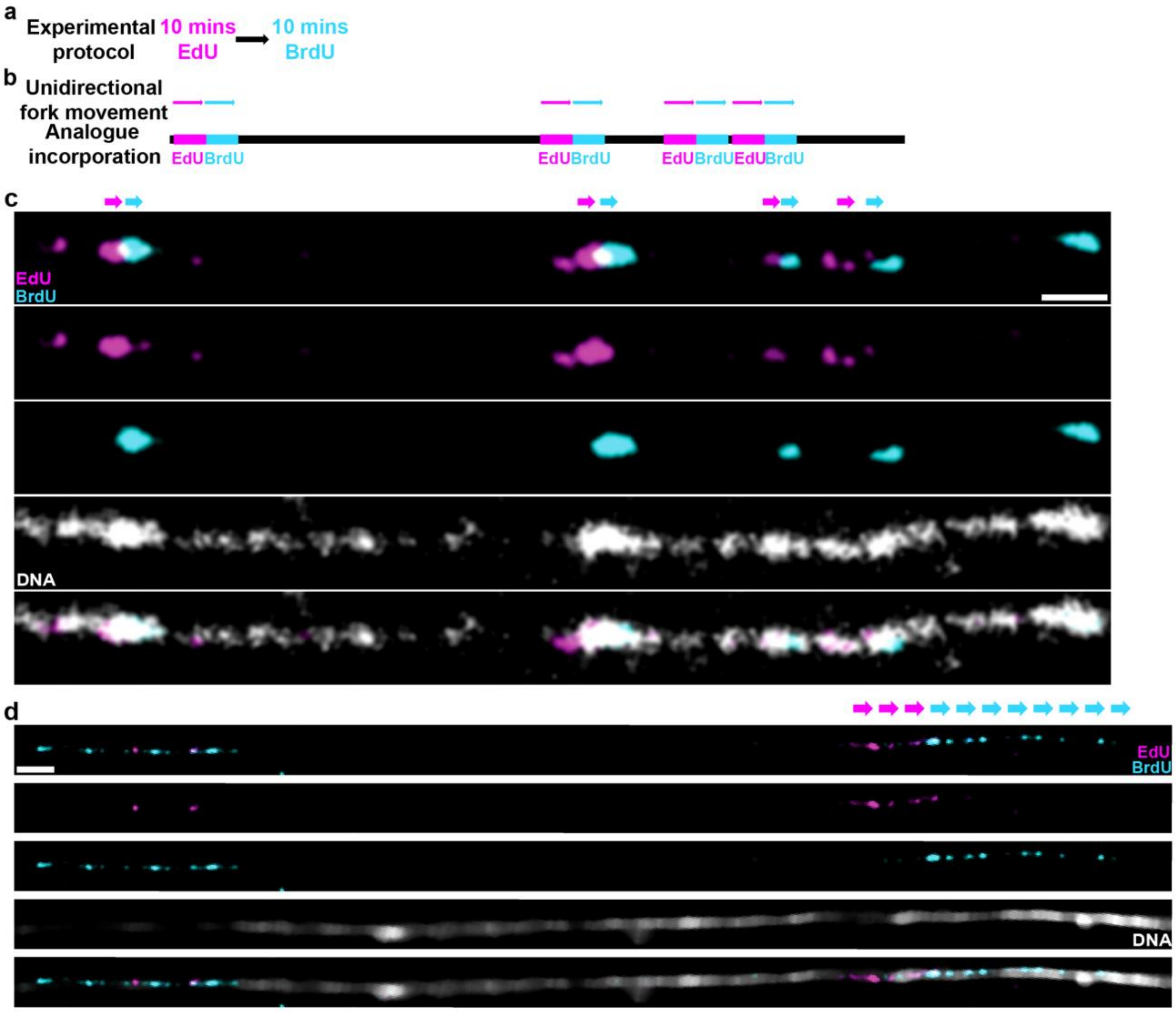

**Supplementary figure 7: DNA label in chromatin fiber and DNA fiber dual-pulse experiments.** (**a**) A cartoon showing experimental protocol. (**b**) Predicted unidirectional fork progression result. (**c**) Unidirectional fork progression pattern from germline-derived chromatin fiber. Multiple replicons show alternation between early label (EdU in magenta) and late label (BrdU in cyan) along one chromatin fiber toward the same direction. DNA label (DAPI) shows continuity between replicons. (**d**) Unidirectional fork progression pattern from germline-derived DNA fiber. DNA label (anti-ssDNA that recognizes all DNA after HCl-treatment) shows continuity between replicons.

**Supplementary Tables:**

**Supplementary table 1: Quantification of histone H4 with imaging on fixed samples.**

| Pair # | Old H4 GSC/GB | New H4 GB/GSC | Pair # | Old H4 SG1/SG2 | New H4 SG2/SG1 |
|---|---|---|---|---|---|
| 1 | 3.95 | 0.58 | 1 | 1.01 | 0.92 |
| 2 | 3.22 | 1.89 | 2 | 1.03 | 0.93 |
| 3 | 3.04 | 0.859 | 3 | 0.85 | 0.82 |
| 4 | 2.95 | 0.98 | 4 | 1.05 | 1.17 |
| 5 | 3.41 | 1.23 | 5 | 1.03 | 0.83 |
| 6 | 5.7 | 1.05 | 6 | 1.31 | 0.77 |
| 7 | 4.86 | 1.19 | 7 | 0.76 | 1.05 |
| 8 | 2.445 | 1.21 | 8 | 1.03 | 1.22 |
| 9 | 1.01 | 1.43 | 9 | 1.069 | 0.98 |
| 10 | 3.15 | 1.038 | 10 | 0.92 | 0.84 |
| 11 | 4.84 | 0.72 | 11 | 1.038 | 1.05 |
| 12 | 5.66 | 0.752 | 12 | 0.89 | 1.22 |
| 13 | 3.21 | 1.75 | 13 | 1.026 | 0.98 |
| 14 | 4.38 | 0.66 | 14 | 0.76 | 0.84 |
| 15 | 5.66 | 1.14 | 15 | 1.04 | 1.05 |
| 16 | 3.21 | 0.887 | 16 | 0.925 | 0.94 |
| 17 | 4.38 | 1.75 | 17 | 1.02 | 0.82 |
| 18 | 1.12 | 0.69 | 18 | 0.76 | 0.95 |
| 19 | 4.31 | 1.15 | 19 | 1.04 | 0.872 |
| 20 | 3.65 | 1.14 | 20 | 0.925 | 1.202 |
| 21 | 0.9 | 0.78 | 21 | 1.02 | 1.147 |
| 22 | 0.76 | 0.57 | 22 | 1.016 | 0.8 |
| 23 | 3.64 | 1.17 | 23 | 0.99 | 1.12 |
| 24 | 3.822 | 0.57 | 24 | 1.00 | 1.03 |
| 25 | 1.337 | 1.17 | 25 | 1.02 | 0.68 |
| 26 | 4.53 | 1.05 | 26 | 1.123 | 1.38 |
| 27 | 2.92 | 1.077 | 27 | 1.01 | 0.93 |
| 28 | 3.08 | 1.03 | | | |
| 29 | 0.94 | 1.287 | | | |
| 30 | 4.87 | 1.149 | | | |
| 31 | 5.97 | 2.116 | | | |
| 32 | 1.39 | 0.815 | | | |
| 33 | 0.98 | 0.982 | | | |

**Supplementary table 2: Quantification of histone H2A with imaging on fixed samples.**

| Pair # | Old H2A GSC/GB | New H2A GB/GSC | Pair # | Old H2A SG1/SG2 | New H2A SG2/SG1 |
|---|---|---|---|---|---|
| 1 | 0.975257657 | 0.98053122 | 1 | 1.088526265 | 1.043770558 |
| 2 | 0.968836869 | 0.966059723 | 2 | 1.016521777 | 1.016185595 |
| 3 | 0.969108816 | 0.941793393 | 3 | 0.94226273 | 0.974142098 |
| 4 | 1.232283465 | 1.633217284 | 4 | 1.067556671 | 0.915380396 |
| 5 | 1.025846378 | 1.041755268 | 5 | 1.009897937 | 0.962468942 |
| 6 | 1.053258093 | 0.999096786 | 6 | 1.042815974 | 0.946732867 |
| 7 | 0.96981108 | 1.186201719 | 7 | 1.019320953 | 0.950287833 |
| 8 | 0.797679181 | 1.357976654 | 8 | 1.222330968 | 0.885426578 |
| 9 | 1.075991617 | 1.122733612 | 9 | 0.995727661 | 1.019966875 |
| 10 | 1.121519519 | 0.901016184 | 10 | 0.930194711 | 1.009292519 |
| 11 | 1.061309268 | 1.103250478 | 11 | 0.931785196 | 0.931550686 |
| 12 | 0.91240285 | 1.066446402 | 12 | 0.986123708 | 0.979875209 |
| 13 | 0.801834862 | 1.204483553 | 13 | 0.925085483 | 1.045018182 |
| 14 | 1.070005651 | 0.987937274 | 14 | 1.019632679 | 1.10359635 |
| 15 | 0.72144534 | 1.247089104 | 15 | 1.035048915 | 1.119932432 |
| 16 | 1.338405425 | 1.246516489 | 16 | 0.969548629 | 0.972565036 |
| 17 | 0.898969072 | 1.154507556 | 17 | 0.972673954 | 1.035350772 |
| 18 | 1.152941753 | 1.019673558 | 18 | 1.023154848 | 1.16412729 |
| 19 | 1.054371002 | 0.823608964 | 19 | 0.978854429 | 1.153198983 |
| 20 | 0.920813893 | 1.027300496 | 20 | 1.055423123 | 1.029364311 |

**Supplementary table 3: Quantification of histone H2B with imaging on fixed samples.**

| Pair # | Old H2B GSC/GB | New H2B GB/GSC | Pair # | Old H2B SG1/SG2 | New H2B SG2/SG1 |
|---|---|---|---|---|---|
| 1 | 0.815283172 | 1.149883726 | 1 | 1.06879687 | 1.036802671 |
| 2 | 0.910809049 | 1.087101455 | 2 | 0.83419399 | 1.01759456 |
| 3 | 0.992565434 | 0.900208136 | 3 | 1.010622669 | 0.861117493 |
| 4 | 0.978807685 | 1.061501775 | 4 | 1.046838138 | 1.052032321 |
| 5 | 1.17406494 | 0.821823967 | 5 | 0.916185819 | 0.874832476 |
| 6 | 0.872269007 | 1.360653409 | 6 | 0.973981549 | 1.326719124 |
| 7 | 1.162729831 | 0.827364081 | 7 | 1.220158888 | 0.749905276 |
| 8 | 1.171490593 | 1.601112878 | 8 | 1.009274627 | 0.968346435 |
| 9 | 1.037883808 | 0.917360074 | 9 | 0.947504382 | 1.170238975 |
| 10 | 0.99783673 | 0.923628319 | 10 | 0.854147825 | 1.174910873 |
| 11 | 0.895076097 | 1.281920327 | 11 | 1.172142501 | 0.858156863 |
| 12 | 0.959500446 | 1.479566305 | 12 | 1.083404453 | 0.845157357 |
| 13 | 1.231086253 | 1.127473807 | 13 | 1.07626037 | 0.923235726 |
| 14 | 0.808704809 | 1.211914894 | 14 | 1.021646558 | 0.920400153 |
| 15 | 1.046689113 | 1.143965517 | 15 | 0.86262317 | 1.172370089 |
| 16 | 0.849082443 | 1.453828829 | 16 | 1.026491198 | 1.939485628 |
| 17 | 1.055365474 | 1.121270452 | 17 | 1.0234375 | 0.972416813 |
| 18 | 1.014446228 | 1.165391969 | 18 | 0.928342031 | 0.856763926 |
| 19 | 0.912596963 | 1.306268241 | 19 | 0.900293686 | 1.918833044 |
| 20 | 1.021970333 | 0.862912736 | 20 | 1.066753078 | 1.144069104 |
| 21 | 1.202621287 | 1.191425723 | 21 | 1.099325769 | 0.662652053 |
| 22 | 0.887726959 | 2.07110666 | 22 | 1.153474545 | 0.59781155 |
| 23 | 0.990796476 | 0.885693395 | 23 | 0.982679645 | 1.374321095 |
| 24 | 0.918245383 | 0.891502847 | 24 | 0.844495944 | 1.205987906 |
| 25 | 0.912221729 | 1.384310526 | 25 | 1.036036036 | 0.852305896 |
| 26 | 0.869454545 | 1.11416998 | 26 | 1.064761181 | 0.948237664 |
| 27 | 1.190698579 | 1.019536742 | 27 | 0.88406336 | 1.059149083 |
| 28 | 0.891282778 | 1.129701061 | 28 | 1.093008455 | 0.998295745 |
| 29 | 1.048793662 | 0.94980315 | 29 | 1.080438985 | 1.300522734 |
| 30 | 1.214413768 | 0.848577475 | 30 | 1.222637781 | 1.0442979 |
| 31 | 0.935564854 | 1.327795976 | 31 | 1.037889226 | 1.10251344 |
| 32 | 0.843987298 | 1.593786228 | 32 | 1.126153435 | 0.977582529 |
| 33 | 1.011382114 | 1.166823456 | 33 | 0.813552882 | 0.91765286 |
| 34 | 0.907251972 | 1.466248278 | 34 | 1.083184342 | 0.968471789 |
| 35 | 0.894008235 | 1.033427164 | 35 | 0.904325323 | 1.031899183 |
| 36 | 1.110156314 | 1.905524681 | 36 | 1.015971606 | 1.013173653 |
| 37 | 0.917364991 | 1.476244026 | | | |
| 38 | 0.964285714 | 1.280842528 | | | |

| | | |
|---|---|---|
| 39 | 0.976632851 | 2.048527984 |
| 40 | 0.96929659 | 0.890530557 |

**Supplementary table 4: Quantification of histone H1 with imaging on fixed samples.**

| Pair# | Old H1 GSC/GB | New H1 GB/GSC |
|---|---|---|
| 1 | 1.231616 | 1.236307 |
| 2 | 1.313828 | 1.371778 |
| 3 | 1.280991 | 1.288979 |
| 4 | 2.686801 | 2.020878 |
| 5 | 1.214697 | 0.676174 |
| 6 | 0.960562 | 0.990234 |
| 7 | 1.147011 | 1.705283 |
| 8 | 1.629132 | 1.252325 |
| 9 | 0.67269 | 0.772935 |
| 10 | 0.846208 | 0.757713 |
| 11 | 1.223122 | 1.269666 |
| 12 | 0.903526 | 0.828135 |

**Supplementary References:**

1. Van Doren, M., Williamson, A.L. & Lehmann, R. Regulation of zygotic gene expression in Drosophila primordial germ cells. *Curr Biol* **8**, 243-6 (1998).
2. Tran, V., Lim, C., Xie, J. & Chen, X. Asymmetric division of Drosophila male germline stem cell shows asymmetric histone distribution. *Science* **338**, 679-82 (2012).
3. Hime, G.R., Brill, J.A. & Fuller, M.T. Assembly of ring canals in the male germ line from structural components of the contractile ring. *J Cell Sci* **109 ( Pt 12)**, 2779-88 (1996).
4. Kolb, H.C., Finn, M.G. & Sharpless, K.B. Click Chemistry: Diverse Chemical Function from a Few Good Reactions. *Angew Chem Int Ed Engl* **40**, 2004-2021 (2001).
5. Moses, J.E. & Moorhouse, A.D. The growing applications of click chemistry. *Chem Soc Rev* **36**, 1249-62 (2007).
6. Koster, D.A., Crut, A., Shuman, S., Bjornsti, M.A. & Dekker, N.H. Cellular strategies for regulating DNA supercoiling: a single-molecule perspective. *Cell* **142**, 519-30 (2010).
7. Wang, J.C. Cellular roles of DNA topoisomerases: a molecular perspective. *Nat Rev Mol Cell Biol* **3**, 430-40 (2002).
8. Kuzminov, A. When DNA Topology Turns Deadly - RNA Polymerases Dig in Their R-Loops to Stand Their Ground: New Positive and Negative (Super)Twists in the Replication-Transcription Conflict. *Trends Genet* **34**, 111-120 (2018).
9. LaMarr, W.A., Yu, L., Nicolaou, K.C. & Dedon, P.C. Supercoiling affects the accessibility of glutathione to DNA-bound molecules: positive supercoiling inhibits calicheamicin-induced DNA damage. *Proc Natl Acad Sci U S A* **95**, 102-7 (1998).
10. Ljungman, M. & Hanawalt, P.C. Localized torsional tension in the DNA of human cells. *Proc Natl Acad Sci U S A* **89**, 6055-9 (1992).
11. Techer, H. et al. Replication dynamics: biases and robustness of DNA fiber analysis. *J Mol Biol* **425**, 4845-55 (2013).
12. Rezaei Poor Kardost, R., Billing, P.A. & Voss, E.W., Jr. Generation and characterization of three murine monoclonal nucleotide binding anti-ssDNA autoantibodies. *Mol Immunol* **19**, 963-72 (1982).
13. Lasko, P.F. & Ashburner, M. Posterior localization of vasa protein correlates with, but is not sufficient for, pole cell development. *Genes Dev* **4**, 905-21 (1990).
